# Molecular and Phenotypic Characterization of Telomere Repeat Binding (TRBs) Proteins in Moss: Evolutionary and Functional Perspectives

**DOI:** 10.1101/2025.06.11.659030

**Authors:** Alžbeta Kusová, Marcela Holá, Ivana Goffová Petrová, Jiří Rudolf, Dagmar Zachová, Jan Skalák, Jan Hejátko, Božena Klodová, Tereza Přerovská, Martin Lyčka, Eva Sýkorová, Yann J. K. Bertrand, Jiří Fajkus, David Honys, Petra Procházková Schrumpfová

## Abstract

Telomere repeat binding (TRB) proteins are plant-specific proteins with a unique domain structure distinct from telomere-binding proteins in animals and yeast. While extensively studied in seed plants, their role in early-diverging plant lineages remain largely unexplored. Here, we investigate TRB proteins in a model moss, *Physcomitrium patens*, to assess their evolutionary conservation and functional significance. Functional analysis using single knockout mutants revealed that individual PpTRB genes are essential for normal development, with mutants exhibiting defects in the two-dimensional (protonemal) stage and, more prominently, in the formation of three-dimensional (gametophore) structures. Some double mutants displayed telomere shortening, a phenotype also observed in TRB-deficient seed plants, indicating a conserved role for TRBs in telomere maintenance. Transcriptome profiling of TRB mutants revealed altered expression of genes associated with transcriptional regulation and stimulus response in protonema. Subcellular localization studies across various plant cell types confirmed that PpTRBs, like their seed plant counterparts, localize prevalently to the plant nucleus and mutually interact. In bryophytes, TRBs form a monophyletic group that mirrors the species phylogeny, whereas in seed plants, TRBs have diversified into two distinct monophyletic groups. Our findings provide the first comprehensive characterization of TRB proteins in non-vascular plants and demonstrate their conserved roles in telomere maintenance, with additional implications for plant development and gene regulation across land plant lineages.

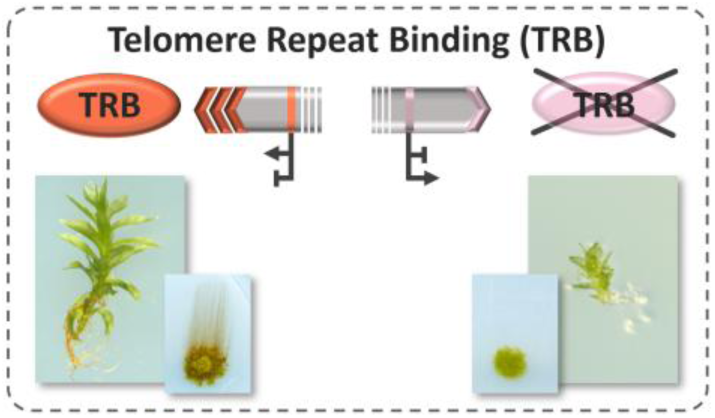

## Introduction

Telomere Repeat Binding (TRB) proteins were initially characterized in seed plants for their roles in telomere maintenance (Marian *et al*., 2003; Schrumpfová *et al*., 2004; Schrumpfová *et al*., 2008) as well as in gene regulation (Schrumpfová *et al*., 2016; Zhou *et al*., 2018; Tan *et al*., 2018; Wang *et al*., 2023; Kusová *et al*., 2023). TRB proteins exhibit a plant-specific domain architecture comprising an N-terminal Myb-like domain, a histone-like (H1/5-like) domain, and a coiled-coil domain at the C-terminus. All five members of the TRB protein family in *A. thaliana* have been shown to bind plant telomeric repeats (TTTAGGG)_n_ through their Myb-like domain (Mozgová *et al*., 2008; Kusová *et al*., 2023). The centrally positioned H1/5-like domain mediates protein-protein interactions, as demonstrated in our previous studies (Kuchar and Fajkus, 2004; Schrumpfová *et al*., 2008).

TRB proteins contribute to telomere maintenance, as demonstrated by telomere length variation in *A. thaliana* mutants in some *AtTRB* genes (Schrumpfová *et al*., 2014; Zhou *et al*., 2018; Amiard *et al*., 2024). All TRB family members in *A. thaliana* are proposed to participate in biogenesis of the enzyme responsible for maintaining telomere length, named telomerase. TRBs interact directly with the catalytic protein subunit of telomerase (Schrumpfová *et al*., 2014; Kusová *et al*., 2023) and with plant homologues of key components of the mammalian telomere complex Shelterin, which is responsible for maintaining telomere sequences (e.g. AtPot1a/b; Protection of Telomeres 1a/b) (Kuchar and Fajkus, 2004; Schrumpfová *et al*., 2008; Kusová *et al*., 2023). In line with that, *in situ* co-localization of AtTRB1 with long telomeric DNA repeats was observed in plant cells (Schrumpfová *et al*., 2014; Dreissig *et al*., 2017).

The Myb-like domain of TRB proteins shares structural similarity with the Myb-like domain of mammalian Telomere Repeat Binding Factors 1 and 2 (hTRF1 and hTRF2) (Broccoli *et al*., 1997), which are the components of the Shelterin complex, responsible for mediating the direct binding of the complex to telomeric DNA sequences. However, unlike TRB proteins, hTRFs possess the Myb-like domain at the C-terminus and lack H1/5-like and coiled-coil domains. Surprisingly, in *Arabidopsis*, similarly structured proteins as mammalian TRFs, named TRF-Like (TRFL) proteins, possess a C-terminal Myb domain but do not participate in telomere maintenance (Fulcher and Riha, 2016) and require an accessory Myb-extension domain for binding telomeric dsDNA *in vitro* (Karamysheva *et al*., 2004; Ko *et al*., 2008).

The function of TRB proteins extends beyond their association with telomeres and telomerase. In addition to binding long terminal telomeric repeats, TRB proteins also interact with other DNA sequences, positioned interstitially, as are short interstitial telomeric sequences, known as *telo*-box motifs (Regad *et al*., 1994; Schrumpfová *et al*., 2016), as well as to Site II motifs (Zhou *et al*., 2018; Wang *et al*., 2023), JMJ14 binding motifs (Wang *et al*., 2023) and other motifs which are present in the promoter regions of numerous genes. The binding of AtTRBs to *telo*-boxes facilitates the recruitment of various protein complexes, such as JMJ14 (Jumonji-domain containing protein 14), PRC2 (Polycomb Repressive Complex 2), or PEAT complex (PWOs-EPCRs-ARIDs-TRBs). These complexes, recruited by TRBs, have diverse roles in regulating gene expression, including histone H3K4 demethylation (Wang *et al*., 2023; Amiard *et al*., 2024) and histone H3K27 methylation (Zhou *et al*., 2018; Kusová *et al*., 2023; Xuan *et al*., 2024), as well as histone H2A deubiquitination and histone H4 acetylation, which may contribute to heterochromatin silencing (Tan *et al*., 2018; Tsuzuki and Wierzbicki, 2018) as well as the formation of an active chromatin state (Zheng *et al*., 2023).

In our previous phylogenetic analysis, we reported evidence of variable rates of TRB gene duplication and copy retention in Streptophyta genomes (Kusová *et al*., 2023). We suggested that TRB proteins first evolved within *Klebsormidiophyceae*; however, the occurrence of TRBs in many early-diverging streptophytes remained unresolved due to the limited genomic data. In Bryophyta we identified TRB genes with very high pairwise sequence similarity to each other.

The genomes of spermatophytes (seed plants) have been shaped by several rounds of whole genome duplication events (WGDs) not shared with Bryophyta and Lycophyta (Clark and Donoghue, 2018). The history of WGD is reflected in the TRB phylogeny as we were able to distinguish two main distinct lineages in seed plants (Kusová *et al*., 2023). In *A. thaliana*, these two TRB lineages are represented by three paralogues (AtTRB1–3), and two additional paralogues (AtTRB4–5), respectively (Schrumpfová *et al*., 2014; Kusová *et al*., 2023).

In the spreading earth moss *Physcomitrium patens* (formerly *Physcomitrella*), three proteins with high sequence similarity, containing a unique Bryophyte-specific motif, were identified through our *in silico* analysis (Kusová *et al*., 2023). The moss *P. patens* was the first bryophyte to have its genome sequenced (Rensing *et al*., 2008; Lang *et al*., 2018) and may serve as model system to investigate the shift from the two-dimensional filamentous phase of the life cycle (protonemata, composed of chloronemal and caulonemal cells) to the three-dimensional (gametophores) stages (Moody, 2022). As in other mosses, the dominance of the haploid phase during the *P. patens* life cycle simplifies genetic manipulations (Prigge and Bezanilla, 2010), and its highly efficient system of homologous recombination (HR) offers a straightforward approach for targeted gene replacement (Schaefer and Zrÿd, 1997; Schaefer, 2002). Altogether, *P. patens* is an excellent system for answering questions in evolutionary developmental biology (reviewed in Rensing *et al*., 2020), including whether TRBs retain their roles and functions, as characterized in *A. thaliana*, in early-diverging plants.

In this study, we conduct a comprehensive characterization of three TRB proteins from the moss *P. patens* (PpTRB1, PpTRB2, and PpTRB3). Phenotypic analysis of various single and double *pptrb* knock-out/knock-down mutants display pronounced morphological growth defects during both the filamentous (protonemata) and shoot-producing (gametophore) stages. Notably, double mutants *pptrb1 pptrb3* and *pptrb2 pptrb3* exhibit telomere shortening which remains consistent across subsequent generations. Transcriptomic analysis of the protonemal stage reveals alterations in the expression of genes encoding proteins involved in DNA transcription regulation, as well as those associated with stimulus response and cellular membrane and peripheral structures. Localization studies in diverse plant cell types demonstrate that PpTRB proteins are predominantly targeted to the nucleus and/or nucleolus. Additionally, mutual interactions between PpTRB proteins were observed in nuclear foci and at the cellular periphery. A comprehensive phylogenetic analysis reveals the presence of *TRB* genes throughout the plant phylogeny, including in streptophyte algae, hornworts, liverworts, mosses and seed plants. Collectively, our findings provide the first detailed characterization of TRB proteins in mosses, demonstrating their roles in regulating plant growth, maintaining telomere integrity, and modulating gene expression. These results highlight the evolutionary conservation of TRB protein functions across early-diverging plant lineages.

## Results

### Mutant in *pptrb* lines exhibit developmental defects

Previously, we have *in silico* identified three *TRB* genes within the *P. patens* genome. These genes exhibit high sequence similarity and encode proteins containing three primary domains: a Myb-like domain, an H1/5-like domain, and a coiled-coil domain. In contrast to the homologues identified in *A. thaliana*, these PpTRB proteins also feature a Bryophyte-specific motif located between the H1/H5 and coiled-coil domains (Kusová *et al*., 2023). In *A. thaliana*, TRBs appear to be functionally redundant, as developmental abnormalities in plants are observed only when mutations affect multiple members of the TRB family (Wang *et al*., 2023; Amiard *et al*., 2024).

To investigate the role of PpTRBs in moss viability, we created various single and double mutant lines (**Supplementary Figure 1** and **Supplementary Figure 2**). The *pptrb1* mutant line was generated by introducing a stop codon into the sequence coding for the Myb-like domain using CRISPR/Cas9-mediated homology-directed repair, resulting in a knock-down mutant line. This was confirmed by RNA-Seq QuantSeq which targets the 3’ end of RNA transcripts (details on QuantSeq are provided below; **Supplementary Figure 3**). The knock-out mutants of *pptrb2* and *pptrb3* were generated using well-established homologous recombination (HR) protocols (Schaefer and Zrÿd, 1997; Kamisugi *et al*., 2006), and also confirmed by QuantSeq. Double mutant lines (*pptrb1 pptrb2*, *pptrb1 pptrb3*, *pptrb2 pptrb3*) were generated from single mutants as described in the Methods section. Despite multiple attempts, we were unable to generate a single *pptrb1* mutant using HR procedures, nor were we able to generate a triple *pptrb1-3* mutant line. Between two and four independently generated mutant lines for each single or double mutant genotype were selected for further analysis (**Supplementary Figure 2**).

The *pptrb2* and *pptrb3* single mutants, as well as all double *pptrb* mutants, exhibited delayed development of macroscopic gametophores compared to the wild-type (WT) moss (**Figure 1a, b; Supplementary Figure 2**). Notably, the double mutants *pptrb1 pptrb3* and *pptrb2 pptrb3* displayed a predominance of protonemal stages over gametophores. Additionally, propidium iodide (PI) staining of 10-day-old protonema revealed that most of the apical caulonemal cells in mutant lines were defective, presenting premature senescence and progressing to cell death (**Figure 1c**).

**Figure 1.**
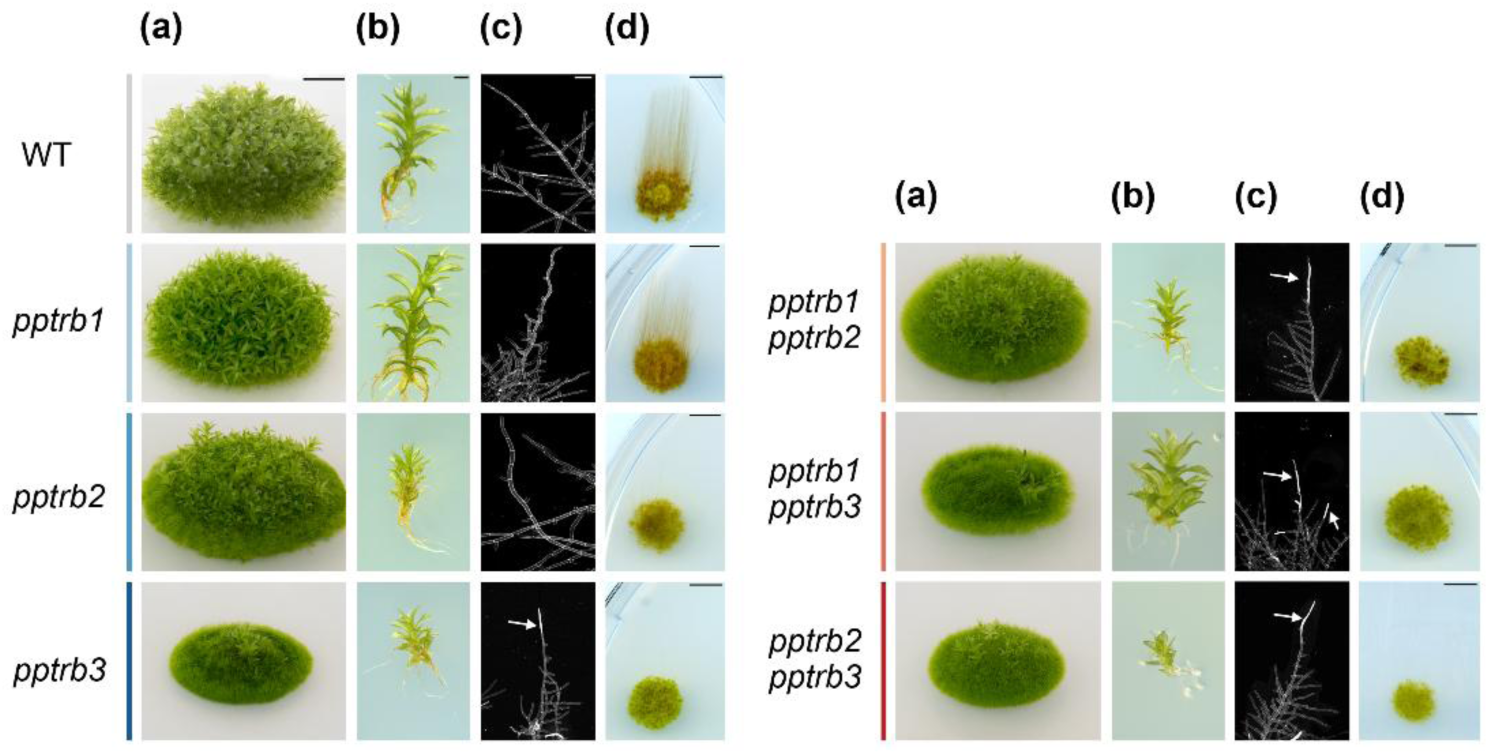
The *pptrb* mutant lines exhibited reduced growth of protonema and, particularly, gametophores. **(a)** Representative morphotypes of 1-month old WT and mutant lines grown on BCDAT medium. Scale bar = 5 mm. **(b)** Close-up view of 1-month-old gametophores of WT and mutant lines. Scale bar = 1 mm. **(c)** Propidium iodide staining of 10-day-old protonema grown BCDAT medium showing defective apical caulonemal cells that are indicated by arrows in *pptrb3*, *pptrb1 pptrb2*, *pptrb1 pptrb3* and *pptrb2 pptrb3* mutant lines. Scale bar = 100 µm. **(d)** Impaired caulonema formation after 3 weeks in the dark conditions. Scale bar = 5 mm.

The transition from chloronema to caulonema in the mutant lines was investigated by exploiting the ability of *P. patens* to develop caulonemata, but not chloronemata, under dark conditions (Cove *et al*., 1978). Protonema grown for one week were transferred – either as homogenized or unhomogenized samples - to new plates, grown for an additional week in the light, and then placed in the dark for three weeks. Under these conditions, WT plants developed long, negatively gravitropic caulonemata, whereas *pptrb2*, *pptrb3*, and all double mutants displayed significantly impaired caulonemal growth (**Figure 1d; Supplementary Figure 4**). This impaired caulonemal formation likely played a critical role in the defects observed during gametophore development, as caulonemata are essential for transitioning to the differentiation of cells that eventually form gametophores (Cove and Knight, 1993; Thelander *et al*., 2018).

Overall, the defects in protonema growth, caulonema development, and gametophore formation detected in the *pptrb* mutants suggest that TRB proteins play a critical role in the early stages of moss development. Specifically, these proteins appear to regulate the transition from the protonemal stage to gametophore formation in *P. patens*. Furthermore, the inability to generate triple *pptrb* mutant lines indicates that PpTRB proteins are likely involved in essential pathways necessary for moss viability and development.

### Role of PpTRB proteins in transcriptional regulation

Since AtTRB proteins are known to be involved in gene regulation in *Arabidopsis* (Schrumpfová *et al*., 2016; Zhou *et al*., 2018; Wang *et al*., 2023), we sought to determine whether PpTRBs retain this role also in *P. patens*. To identify genes differentially regulated by PpTRBs, we performed QuantSeq targeting 3’ end of the polyadenylated RNA isolated from 7-day-old protonema from single and double *pptrb* mutants and compared the results to WT plants.

QuantSeq analysis revealed distinct differences in gene expression profiles between individual single and double *pptrb* mutants, analyzed across three biological replicates. Notably, mutants with *pptrb3* (either single or double), are clearly separated from other genotypes in the Principal Component Analysis (PCA) plot (**Figure 2a**). Numerous differentially expressed genes (DEGs), both up- and down-regulated, were identified in *pptrb* mutant plants compared to the WT (**Supplementary Table 1**).

**Figure 2.**
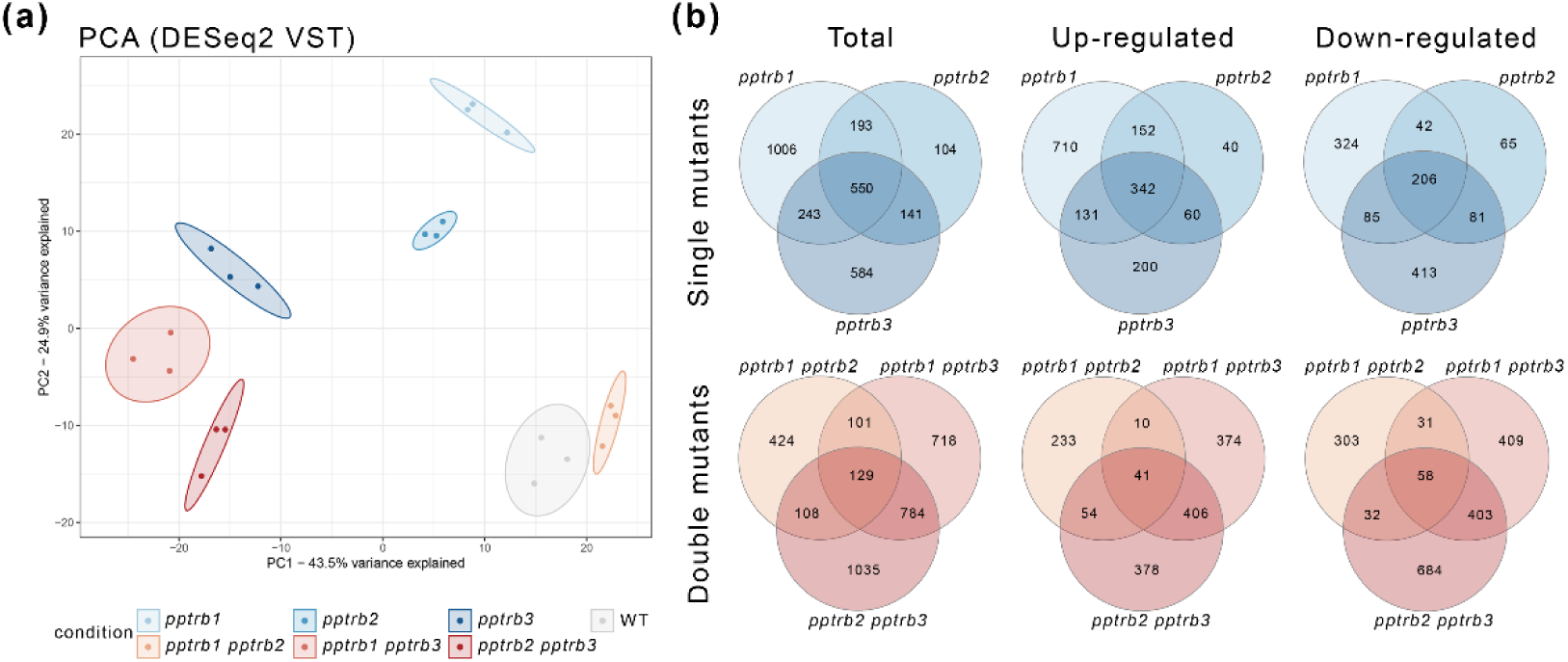
Differential gene expression in *pptrb* mutants. **(a)** Principal component analysis (PCA) based on DESeq2 with variance-stabilizing transformation (VST) reveals transcriptomic differences among *pptrb* single and double mutants. Each mutant line is represented by a distinct color. **(b)** Venn diagram illustrating the overlap of differentially expressed genes (DEGs) among *pptrb* single and double mutants. The total number of DEGs, as well as the subsets of up-regulated and down-regulated genes, are indicated.

Overlapping DEGs among the mutants that were significantly up- or down-regulated are depicted in Venn diagrams (**Figure 2b**). A total of 550 and 129 DEGs were found to overlap among single and double mutant datasets, respectively. Interestingly, the overlap between *pptrb1 pptrb3* and *pptrb2 pptrb3* was higher than the overlap between *pptrb1 pptrb3* and *pptrb1 pptrb2* (**Supplementary Figure 5; Supplementary Figure 6**).

Gene Ontology (GO) analysis of RNA-seq data for up-regulated DEGs revealed significant enrichment in GO Biological Process (GO:BP) terms ‘regulation of DNA-templated transcription’, ‘response to stimuli’. In GO Cellular Components (GO:CC), enriched terms included ‘cell periphery’, ‘membrane’, ‘cell wall’ and ‘extracellular region’. For down-regulated DEGs, GO:BP analysis highlighted terms related to ‘compound of biosynthetic processes’, ‘regulation of DNA-templated transcription’. The GO:CC analysis of the down-regulated DEGs showed enrichment in ‘cell periphery’ and ‘plasma membrane’ terms (**Figure 3 and Supplementary Figure 7**).

**Figure 3.**
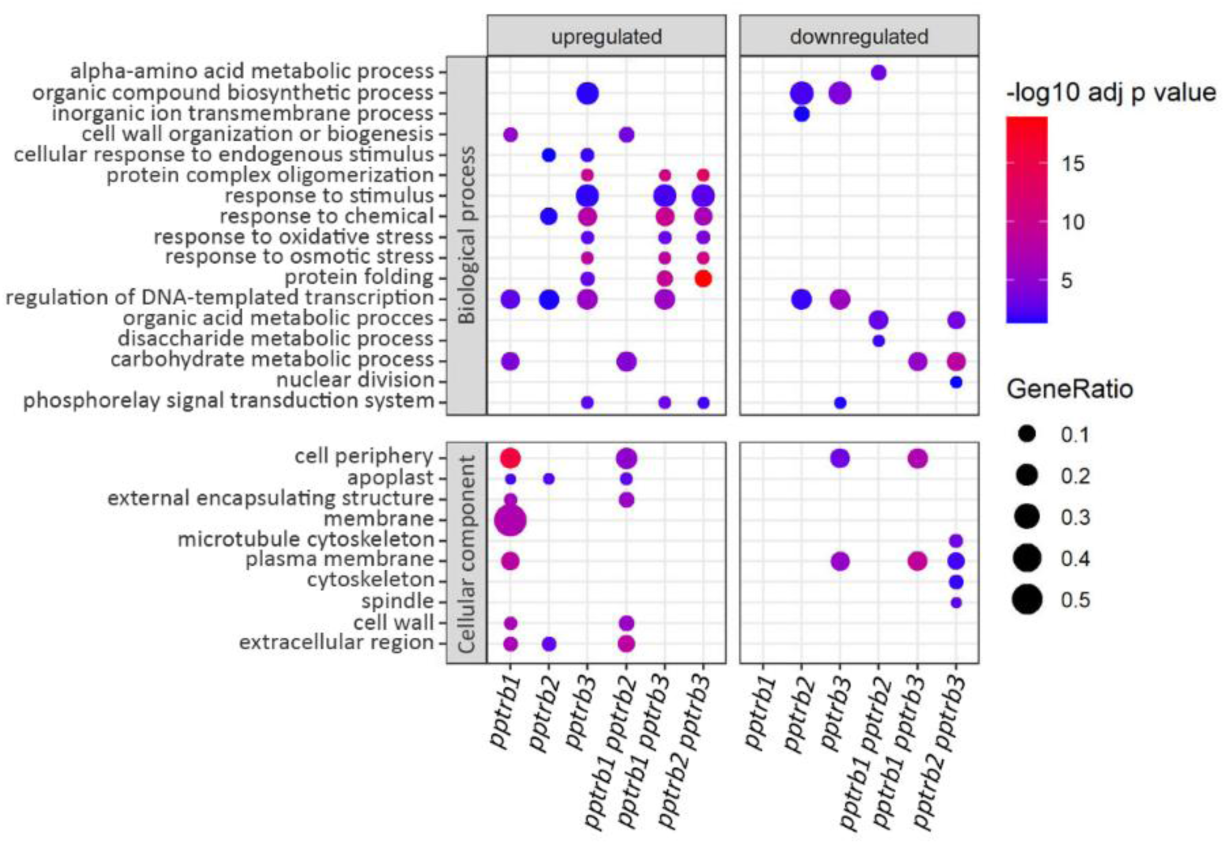
Gene Ontology (GO) enrichment analysis of differentially expressed genes in *pptrb* mutant lines. GO enrichment analysis was performed separately for upregulated and downregulated genes identified in *pptrb* mutants relative to WT. Among the upregulated DEGs, GO terms significantly enriched in the Biological Process (GO:BP) category included ‘regulation of DNA-templated transcription’ and ‘response to stimuli’. Within the Cellular Component (GO:CC) category, enriched terms were associated with ‘cell periphery’, ‘membrane’, ‘cell wall’, and ‘extracellular region’. In downregulated DEGs showed GO:BP enrichment in ‘compound biosynthetic processes’ and again in ‘regulation of DNA-templated transcription’. GO:CC terms for these genes were similarly enriched in ‘cell periphery’ and ‘plasma membrane’. Adjusted p-values for each GO term were derived from g:Profiler. GeneRatio – the ratio of DEGs annotated to the GO term calculated by dividing intersection_size by query_size from the g:Profiler. For full GO enrichment analysis, see **Supplementary Figure 7**.

We utilized our custom program GOLEM (Gene RegulatOry ELeMents) (Nevosád *et al*., 2025), to analyze the frequency and distribution of *telo*-box motifs, a key recognition motif for TRB proteins within gene promoters relative to the transcription start site (TSS). The analysis was conducted on genes with the highest transcription levels in single and double *pptrb* mutant lines, comparing them to both the genome-wide distribution and the frequency in wild-type (WT) plants. The *telo*-box motif (Schrumpfová *et al*., 2016) exhibited an increased frequency of occurrence in the promoters of highly expressed genes in *pptrb* mutants compared to WT plants. Genes containing a *telo*-box motif near their TSS accounted for approximately 15-27% of the most highly transcribed genes in *pptrb* mutant lines but to only 9% in WT plants. Additionally, in contrast to the more dispersed genome-wide distribution, which reflects motif presence regardless of gene transcription, the *telo*-box motif was clearly localized upstream of the TSS in genes highly transcribed in protonema. In contrast, other motifs, such as JMJ14 and Site II (both previously linked to AtTRB proteins; (Zhou *et al*., 2018; Wang *et al*., 2023)), did not show a similar enrichment relative to TSS (**Supplementary Figure 8**).

To further explore these genes, we performed GO analysis on the targets identified by the GOLEM program, that exhibited both the highest transcription levels in *pptrb* mutant plants and the presence of a *telo*-box in their promoters (±100 bp from the TSS). This analysis revealed a higher number of genes coding for ribosomal proteins in the mutant lines compared to the wild type (**Supplementary Table 2**). This finding is consistent with previous observations of Procházková Schrumpfová et al. (Schrumpfová *et al*., 2016), who demonstrated via ChIP-Seq that AtTRB1 binds to *telo*-box motifs in the promoters of genes associated with translation machinery, particularly ribosomal protein genes. Interestingly, when we analyzed only the promoters of DEGs, irrespective of their transcription levels, no significant enrichment of motifs or prevalence of ribosomal genes was observed.

Altogether, our RNA-seq analysis of *pptrb* single and double mutants revealed distinct expression profiles, particularly in mutants involving *pptrb3*. GO analysis suggested that TRB are involved in regulating genes associated with transcriptional regulation, response to various stimuli and cellular periphery functions. Furthermore, the *pptrb* mutants exhibited higher transcript levels of genes containing *telo*-box motifs near the TSS, particularly those encoding ribosomal proteins.

### Loss of PpTRBs results in telomere shortening

To determine whether the function of plant-specific TRB proteins in telomere maintenance is conserved in mosses, as observed in *attrb1-3* (Schrumpfová *et al*., 2014; Zhou *et al*., 2018) but not *attrb4-5* (Amiard *et al*., 2024), telomere lengths were analyzed in *P. patens* mutants lacking TRB proteins.

The terminal restriction fragments (TRF) analysis was performed on DNA isolated from WT and *pptrb* single and double mutant lines across three consecutive passages of protonemas, beginning with the third passage. This design allowed us to investigate whether possible telomere shortening progressed over time, as previously observed, for example, in *attrb1* mutants (Schrumpfová *et al*., 2014).

Genomic DNA was digested by *Mse*I restriction enzyme, and intact telomeric fragments were visualized via Southern hybridization with radioactively labeled telomeric probes (**Figure 4a**). The median telomere length was quantified using Web-based Analyzer of the Length of Telomeres (WALTER) toolset (Lyčka *et al*., 2021). Telomere shortening was observed in two double mutant lines, *pptrb1 pptrb3* and *pptrb2 pptrb3*, both of which contain mutation in the *PpTRB3* gene (**Figure 4b**). The median telomere length in these mutant lines was approximately 650 bp, roughly half the length of a WT plant telomere, which exhibited a median length of approximately 1250 bp. Interestingly, this telomere shortening appeared to occur prior to the first analyzed passage and did not progress further across subsequent passages of protonema. In contrast, no changes in the telomere lengths were observed in single *pptrb1* and *pptrb2* mutant lines, while slight changes were observed in *pptrb3* mutant line. Altogether, the observed telomere shortening in several *pptrb* mutant lines indicates that TRB proteins play a critical role in telomere maintenance in mosses, suggesting a conserved ancestral function present in the last common ancestor of tracheophytes (vascular plants) and bryophytes (non-vascular plants) that diverged during the Cambrian, 515–494 million years ago (Harris *et al*., 2022).

**Figure 4.**
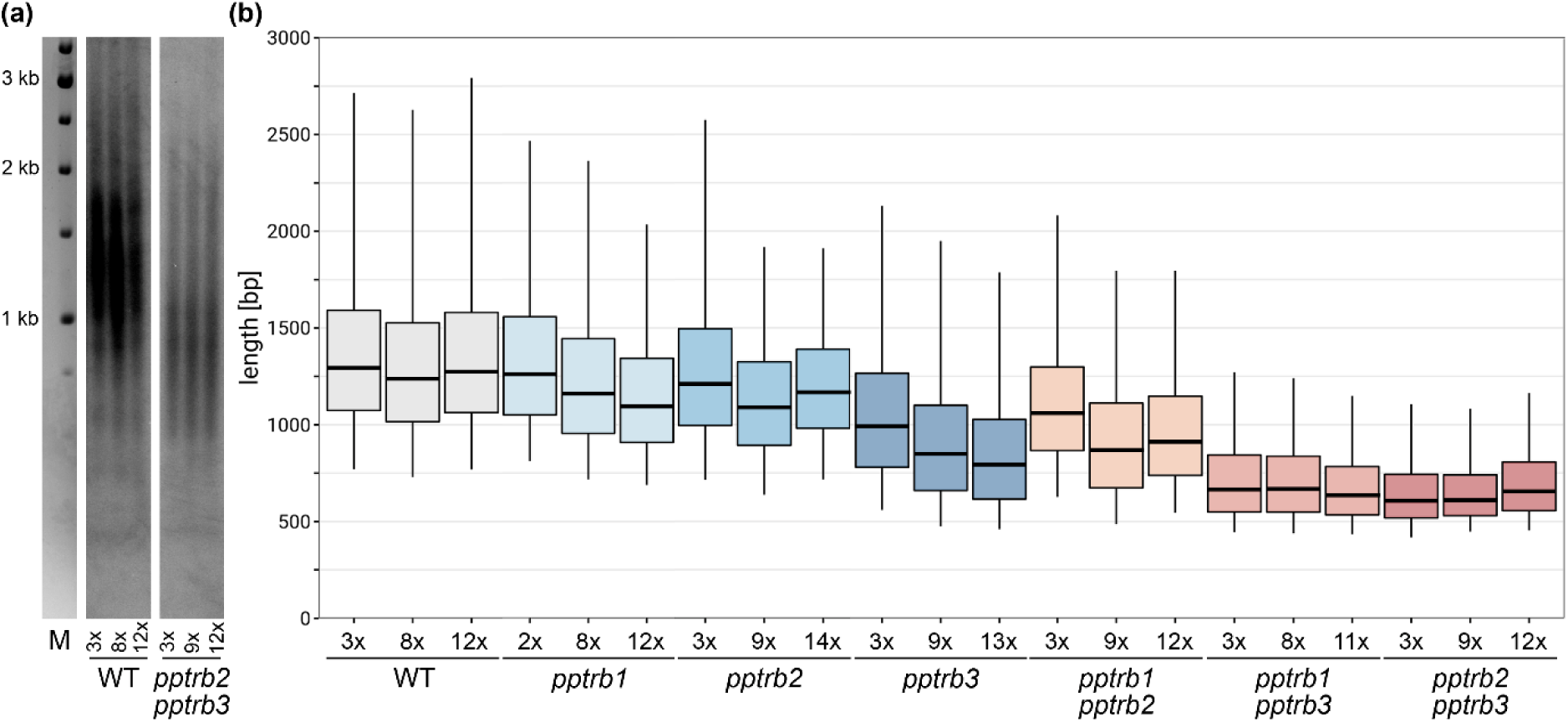
Telomere length analysis reveals stable telomere shortening in some of *pptrb* mutant lines. **(a)** Telomere Restriction Fragment (TRF) Southern blot analysis of three biological replicates of *P. patens* WT and *pptrb2 pptrb3* mutants across the indicated protonemal passages. Molecular weight DNA markers (in kb) are shown. **(b)** Boxplots represent mean TRF length distributions across three analyzed passages. The graphical representation was generated using WALTER (Lyčka *et al*., 2021). Shortened telomeres were detected predominantly in the *pptrb1 pptrb3* and *pptrb2 pptrb3* double mutant lines, with median length of approximately 600 bp.

### PpTRB proteins show conserved nuclear and nucleolar localization

To determine the subcellular localization of TRB proteins in mosses, we generated translational fusions of PpTRB1, 2 and 3 proteins with green fluorescent protein (GFP). These constructs were placed under the control of the constitutive CaMV 35S promoter (35S::GFP:PpTRB1-3) (Karimi *et al*., 2002). Using polyethylene glycol (PEG)-mediated transient transformation of *P. patens* protoplasts derived from 7-day-old protonema cells, we observed that PpTRBs preferentially localized to the nucleus and nucleolus (**Figure 5a**). Specifically, PpTRB1 and PpTRB2 proteins are localized within both the nucleus and nucleolus, whereas PpTRB3 was exclusively localized to the nucleolus. This subcellular distribution in *P. patens* was consistent with the localization of AtTRB1-3 proteins in *Arabidopsis* cells (Dvořáčková *et al*., 2010) and the localization of AtTRB1-4, but not AtTRB5, in tobacco leaf epidermal cells (Zhou *et al*., 2016; Kusová *et al*., 2023).

**Figure 5.**
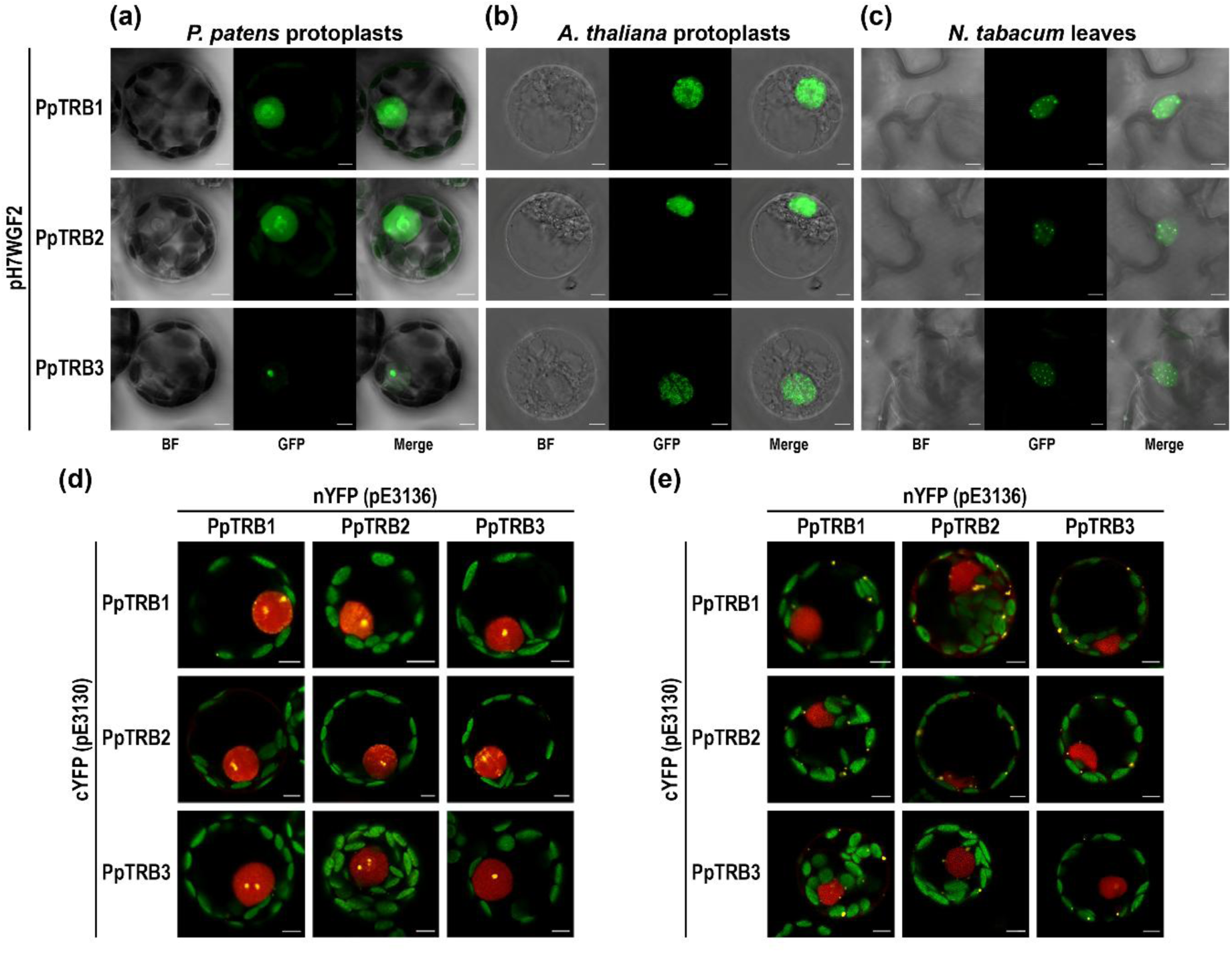
PpTRBs localize to the nucleus across different plant cell types from various organisms and exhibit mutual interactions. (a–c) Subcellular localization of PpTRB1–3 proteins fused to GFP, transiently expressed in: (a) *P. patens* protoplasts derived from 7-day-old protonema, (b) *A. thaliana* protoplasts derived from seedling roots, (c) *N. tabacum* (SR1) leaf epidermal cells. In all examined cell types, PpTRB proteins localize predominantly to the nucleus, with minor variations observed between species. Merged channels of Bright Field (BF) and GFP are shown. Scale bars = 5 µm. **(d, e)** Protein–protein interactions among PpTRBs, fused with nYFP or cYFP fragments, were detected using Bimolecular Fluorescence Complementation (BiFC) in *P. patens* protonemal protoplasts. Reconstituted YFP fluorescence (protein interaction) is shown together with mRFP fluorescence (internal control for transformation and expression) as merged confocal images: **(d)** fluorescence foci of varying numbers and sizes are localized primarily in the nucleus, **(e)** in some protoplasts, fluorescence foci are also observed in the cytoplasm at the cellular periphery. For individual fluorescence channels corresponding to Fig. 5d, e, see Supplementary Figure 9 and 10, respectively. Scale bars = 5 µm.

To further investigate the subcellular localization, we expressed PpTRB-GFP constructs in *Arabidopsis thaliana* and *Nicotiana tabacum* cells. PEG-mediated transient transformation of *A. thaliana* protoplasts, derived from seedling roots, revealed that PpTRBs predominantly localized to the nucleus, though with a more dispersed pattern compared to *P. patens* protoplasts (**Figure 5b**). In contrast, transient *Agrobacterium*-mediated transformation of *N. tabacum* (cv. Petit Havana SR1; referred to hereafter as SR1) leaf epidermal cells also showed nuclear localization of PpTRBs, albeit with a slightly different distribution compared to both *P. patens* and *A. thaliana* protoplasts (**Figure 5c**).

Overall, the PpTRBs demonstrate preferential nuclear localization being conserved in *Physcomitrium, Arabidopsis* and *Nicotiana* with only subtle species-specific variations. Notably, the localization pattern of PpTRBs in *P. patens* protoplasts resembles that of AtTRB1-3 in *Arabidopsi*s cells (Dvořáčková *et al*., 2010).

### PpTRB mutually interact in the nucleus and/or nucleolus, as well as at cellular periphery

To investigate whether the mutual interactions observed among all five AtTRB proteins (Mozgová *et al*., 2008; Kusová *et al*., 2023) are conserved also in moss, we employed a Bimolecular Fluorescence Complementation (BiFC) assay. The coding sequences of *PpTRB1-3* genes were fused to sequences coding for N- and/or C-terminal fragments of yellow fluorescent protein (nYFP and cYFP). Protoplasts derived from 7-day-old protonema cells were co-transformed with mRFP−VirD2NLS construct, which marks the nucleus (Citovsky *et al*., 2006). Our observations revealed that all three PpTRB proteins form homo- and hetero-dimers, as evidenced by nucleoplasmic fluorescence foci of varying numbers and sizes localized predominantly in the nucleus (**Figure 5d**, **Supplementary Figure 9**). Interestingly, fluorescence foci were also observed at the cellular periphery in approximately one-third of the analyzed protoplasts (**Figure 5e**, **Supplementary Figure 10).**

To further validate the mutual interactions of PpTRBs in plant cells, PpTRBs were fused with either green or red fluorescent proteins (GFP, RFP). Due to the challenge of measuring fluorescence lifetime in partially swirling protoplasts in solution, these analyses were performed on transiently transformed *N. tabacum* SR1 leaf epidermal cells. Co-localization of GFP and RFP signals implied possible interaction of PpTRB1-3 proteins in nucleoplasmic foci of varying numbers and sizes (**Figure 6a-c, Supplementary Figure 11**). The specific protein-protein interactions at subcellular precision were confirmed using Fluorescence Lifetime Microscopy - Förster Resonance Energy Transfer (FLIM-FRET; **Figure 6d-f**).

**Figure 6.**
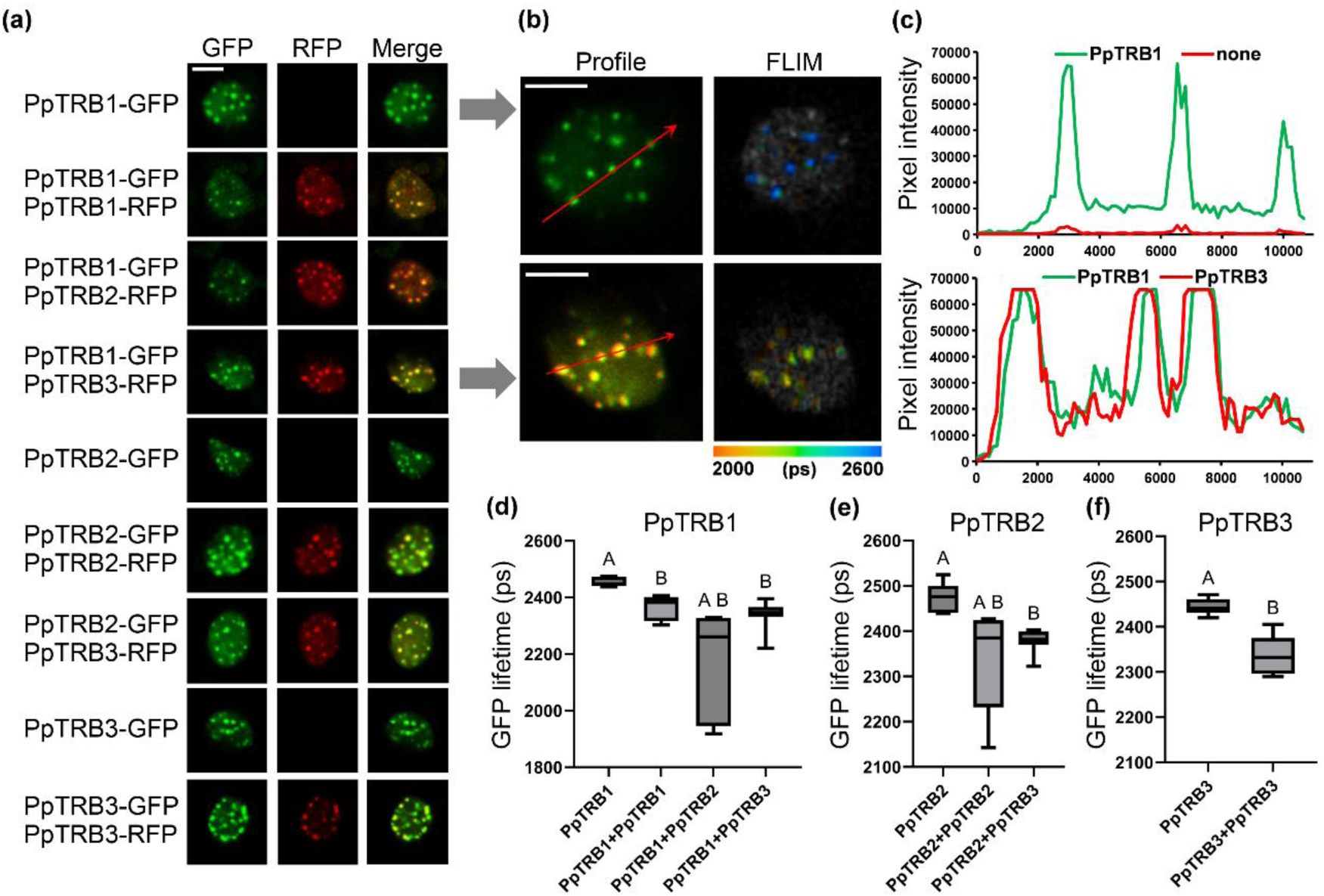
PpTRBs nuclear speckles locate and interact at same spots in nucleoplasm. **(a)** Representative confocal images of subnuclear localization of PpTRBs-GFP/RFP with merged images. **(b)** PpTRB1-GFP and PpTRB3-RFP form nuclear speckles and **(c)** colocalizes at the same foci; scale bars = 5 µm. **(c)** Intensity plots for regions of interest (red arrows) indicate co-localization of PpTRB1 with PpTRB3 in nuclear speckles. For intensity plots of individual samples see Supplementary Figure 10. **(d-f)** GFP fluorescence lifetime measured in a FLIM-FRET interaction assay using the indicated vector combinations transiently expressed in *Nicotiana tabacum* leaves. Error bars represent means ± SD; n = 10, and letters indicate statistical significance (two-way ANOVA and Tukey’s post hoc test).

Overall, our analyses demonstrate that PpTRB proteins in *P. patens* exhibit mutual interactions similar to those observed in AtTRB proteins. The interactions of PpTRBs are predominantly nuclear and localized in nucleoplasmic foci, resembling the behavior of AtTRB1-4. However, similar to AtTRB5, whose homodimeric interactions are predominantly cytoplasmic (Kusová *et al*., 2023), PpTRBs also exhibit interactions in the cytoplasm, particularly near the cellular periphery, in a subset of *P. patens* protoplasts.

### Phylogeny of plant TRB proteins

Although we previously identified a single plant-specific member of the TRB family in *Klebsormidium nitens* (Klebsormidiophyceae) (Kusová et al 2023), TRB presence in other classes of streptophyte algae (Mesostigmatophyceae, Chlorokybophyceae, Charophyceae, Charophycae, Coleochaetophyceae), hornworts or liverworts within Bryophyta, remained unresolved. To trace the evolutionary history of TRB proteins in Streptophyta, we surveyed for TRB homologs across fully sequenced and annotated genomes, a proteome, as well as newly available transcriptomes, representing key groups within Streptophyta.

We probed 11 annotated genomes, proteome and 21 previously assembled transcriptomes. We further assembled 30 transcriptomes from publicly available data. Genomes and transcriptomes were evaluated for BUSCO sequences completeness. Our dataset consisted of 444 sequences (see **Supplementary Table 3**). In several instances one or several of the three domains defining TRBs (H1/5-like, MYB, coiled-coil) were missing. The coiled-coil motif seems to be the most diverse at the sequence level, showing large variations and clade specific features. When probing reference proteomes, we identified, in several instances, a full length TRB associated with a truncated copy within the same species (e.g., *Nitella flexilis*), which could indicate pseudogenization. However, the presence of truncated versions could in some cases be ascribed to genome annotation errors as we were able to obtain transcripts of the canonical architecture in closely related species for these duplicated genes. Within the most basal lineages of Streptophytes, TRBs were observed in Chlorokybophyceae, but we failed to detect them in the sister clade of Mesostigmatophyceae (**Figure 7a**). We successfully retrieved TRBs from four out of five cryptic species of *Chlorokybus.* Here, the TRB structure was unusual, showing a disordered C-terminal coiled-coil motif.

**Figure 7.**
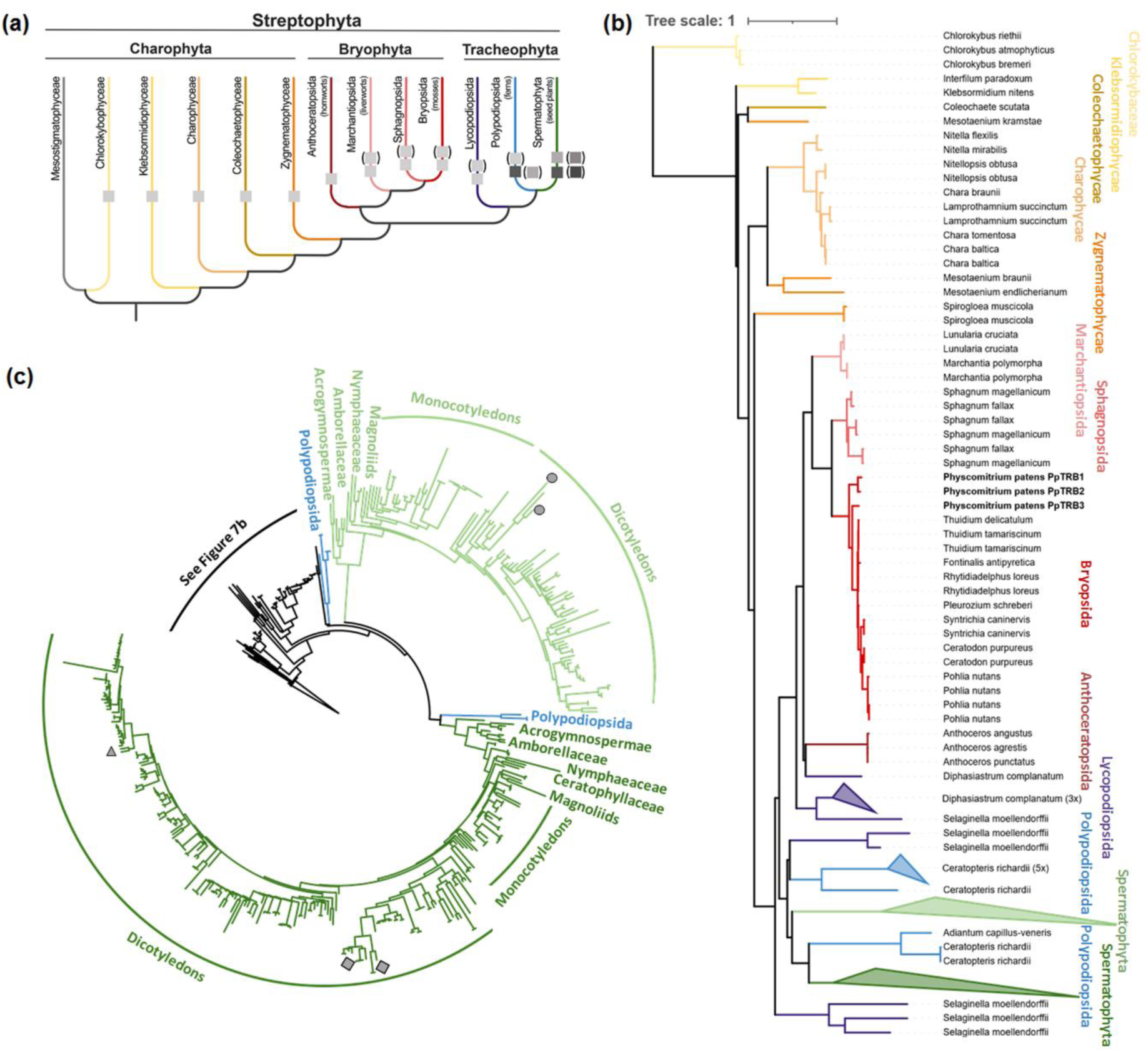
Phylogenetic analysis of TRB proteins across Streptophyta. **(a)** A simplified evolutionary overview illustrating the major lineages within Streptophyta containing identified TRB proteins. The emergence of a single TRB protein is first observed in streptophyte algae, specifically within the Chlorokybophyceae. Subsequent whole-genome duplication (WGD) events contributed to the diversification of TRB proteins into several homologous variants in mosses and lycophytes. In Spermatophyta (seed plants), further diversification resulted in the presence of multiple distinct TRB homologues, which cluster into two major evolutionary lineages. The evolutionary framework is adapted from Rensing (2020) and Cheng *et al*. (2019). **(b)** Distribution of TRB proteins across various streptophyte taxa, with a particular focus on streptophyte algae and bryophytes. In bryophyte lineages such as Marchantiopsida, Bryopsida, and Sphagnopsida, TRB proteins form a monophyletic group. In contrast, TRB proteins in Spermatophyta exhibit greater diversification. **(c)** In Spermatophyta, TRB proteins have diverged into two distinct clades, indicated by light and dark green branches. The *A. thaliana* paralogs AtTRB1 (triangle), AtTRB2, and AtTRB3 (squares) are grouped within the dark green branch, while AtTRB4 and AtTRB5 (circles) belong to the light green branch (Kusová *et al*., 2023).

The TRB phylogeny demonstrated a strong departure from the established streptophyte topology especially in charophytes. Although, both Klebsormidiophyceae and Charophyceae were resolved as monophyletic (**Figure 7b, Supplementary Figure 12**), we recovered Zygnematophyceae as polyphyletic. Withing Zygnematophytes, we failed to get a positive match in several representatives within the Desmidiales order. It is noteworthy that Desmidiales were characterized by a large proportion of missing BUSCO genes (**Supplementary table 3**) which might indicate a severe genome contraction leading to the loss of *TRB*. Moreover, *Spirogloea muscicola* branched in proximity to embryophytes, fitting the recent hypothesis of a close relationship between these two groups and the species divergence from other members of the Zygnematophyceae (Cheng *et al*., 2019; Hess *et al*., 2022).

Within bryophytes, Marchantiopsida (liverworts), Bryopsida (joint-toothed mosses), Sphagnopsida (peat mosses), each formed a monophyletic group with several independent duplications located within them. Anthocerotopsida (hornworts) was positioned as bryotphytes’ sister clade as expected (One Thousand Plant Transcriptomes Initiative 2019 (Leebens-Mack *et al*., 2019)). The evolutionary patterns were more complex within Lycopodiopsida (lycophytes) where a total of 6 putative TRBs from *Selaginella moellendorffii* branched at three distinct positions: at the base of embryophytes, sister to bryophytes and sister to Polypodiopsida (ferns). However, the other representative of lycophytes in our dataset, *Diphasiastrum complanatum,* possessed only TRBs which were associated with bryophyte TRBs. The unexpected positions of *S. moellendorffii* paralogs could be explained by common phylogenetic reconstruction artifacts (e.g., long branch attraction, model misspecification) or they could hint at a more complex evolutionary history of *TRB*s including horizontal gene transfer or gene loss in other clades.

Our phylogeny further demonstrated that a *TRB* duplication occurred before the divergence of Spermatophyta, producing one lineage containing the *AtTRB1-3* paralogs and the other one with the *AtTRB4-5* paralogs (see Kusová *et al*., 2023) **(Figure 7c; Supplementary Figure 12)**. The established sister clade relationship between Acrogymnospermae and Angiospermae was recovered. The topology of the Acrogymnospermae subtree suggested a gene duplication within the Pinopsida family. The relationships within Monocots and Eudicots were generally consistent with the accepted phylogeny. We confirmed the Eudicots’ gene duplication within the lineage containing the *AtTRB1-3* paralogs. Despite sharing all its nested WGDs with closely related species (e.g., *A. lyrata* and *A. arenosa*), *A. thaliana* was the only species in our dataset containing the full complement of TRB copies.

In summary, our analysis indicates that TRB proteins originated in Chlorokybophyceae and are present across streptophyte algae. In bryophytes, TRBs form a monophyletic group, whereas in seed plants they diversified into two distinct groups following an early duplication event.

## Discussion

In contrast to animals and yeast, plants have evolved a unique family of proteins known as TRBs, characterized by a distinct domain architecture comprising an N-terminal Myb-like domain, an H1/5-like domain, and a C-terminal coiled-coil domain. In seed plants, TRB proteins play a pivotal roles, particularly in structure and maintenance of telomeres, and regulation of gene expression via epigenetic complexes associated with gene regulatory elements (Schrumpfová *et al*., 2004; Schrumpfová *et al*., 2016; Zhou *et al*., 2016; Kusová *et al*., 2023).

To date, the function of TRB proteins have been extensively characterized only in seed plants, particularly in *A. thaliana* (Schrumpfová *et al*., 2004; Schrumpfová *et al*., 2008; Schrumpfová *et al*., 2014; Schrumpfová *et al*., 2016; Dreissig *et al*., 2017; Zhou *et al*., 2018; Kusová *et al*., 2023; Teano *et al*., 2023; Wang *et al*., 2023; Amiard *et al*., 2024) and partially in apple (*Malus domestica*) (An *et al*., 2021) and rice (Xuan *et al*., 2024). To fill in missing knowledge on these proteins in earlier branching land plants, we characterized here TRB proteins by examining their function in *P. patens*.

### Impact of *pptrb* mutations on moss development

One of the key evolutionary innovations in land plants was the shift from two-dimensional (2D) to three-dimensional (3D) growth (Moody, 2022). This 3D growth is a hallmark feature of land plants. In the moss *P. patens*, a representative of the earliest lineage of land plants, 3D growth (gametophore) is preceded by an extended 2D filamentous growth (protonema) phase, although the former is not a prerequisite for survival in *P. patens* (Moody, 2019).

In this study, we demonstrate that in the moss *P. patens*, three highly similar TRB proteins are transcribed during the 2D protonemal stage. In contrast to mosses, seed plants predominantly exhibit two main diverging lineages of TRBs (Kusová *et al*., 2023). In the seed plant *A. thaliana*, five paralogues have been identified across these two lineages: AtTRB1, AtTRB2, and AtTRB3 in one lineage, and AtTRB4 and AtTRB5 in the other (Schrumpfová *et al*., 2014; Kusová *et al*., 2023). Growth or developmental phenotypes are only observed in mutants lacking all members of a single lineage (Wang *et al*., 2023; Amiard *et al*., 2024).

Our findings suggest that in *P. patens*, TRB proteins are essential for cell viability, as we were unable to generate a plant mutant for all three PpTRB proteins. Even single knockouts for *PpTRB2* and *PpTRB3* show significant phenotypic changes, including reduced 3D stage and gametophore formation. We were unable to achieve a complete knockout of *PpTRB1* using HR, as only knockdown lines generated via CRISPR-Cas were obtained. In *pptrb3* mutants and all double mutant lines, the apical caulonemal cells, which play a crucial role in the transition to differentiated cells forming gametophores, were frequently defective, presenting premature senescence and progressing to cell death.

Whether the essentiality of TRB proteins is linked to their role in early moss development, particularly in regulating the transition from 2D protonemal stages to 3D gametophore formation, remains to be further investigated.

### Functional conservation and divergence of PpTRBs

TRB proteins were first identified in maize and *A. thaliana* for their ability to bind telomeric DNA (Marian *et al*., 2003; Schrumpfová *et al*., 2004). This role was confirmed by telomere shortening in *Arabidopsis* mutants lacking *AtTRB1–3*, but not *AtTRB4–5* (Schrumpfová *et al*., 2014; Lee and Cho, 2016; Amiard *et al*., 2024). In addition to binding telomeric DNA, A. thaliana TRBs also associate with promoter regions of various genes containing *telo*-boxes, Site II motifs, or JMJ14 binding motifs, likely contributing to epigenetic regulation (Schrumpfová *et al*., 2016; Zhou *et al*., 2018; Wang *et al*., 2023). Our findings indicate that TRB proteins are involved in telomere maintenance not only in flowering plants but also in mosses. Telomere shortening was observed in *pptrb* mutant lines, specifically those with mutations in the *PpTRB3* gene (*pptrb3*, *pptrb1 pptrb3* and *pptrb2 pptrb3*). These findings are consistent with findings in *Arabidopsis* plants mutants lacking AtTRB1 (Schrumpfová *et al*., 2014) and AtTRB1-3 (Zhou *et al*., 2018). The significance of these findings lies in the fact that TRB proteins in *P. patens* differ from the two distinct groups of TRBs characterized in seed plants. Telomere shortening, indicative of chromosome instability, was also associated with morphological changes in the *P. patens* mutant lines. The absence of telomere shortening in single mutants suggests possible functional redundancy among the three highly similar TRB proteins in *P. patens*.

To elucidate the molecular mechanisms underlying the observed phenotypic changes, we conducted a differential expression analysis using RNA-seq data. The identification of differentially expressed genes (DEGs) in these mutants provided insights into the cellular pathways affected by TRB protein loss. Notably, several genes related to regulation of DNA-templated transcription or various stress responses were significantly altered, highlighting the diverse roles TRB proteins play in maintaining cellular homeostasis. Interestingly, the *pptrb* mutants exhibited higher transcription levels of genes containing *telo*-box motifs near TSS, particularly those encoding ribosomal proteins. In other analyzed motifs already described as to be recognized by TRB proteins as Site II (TGGGCY) (Hervé *et al*., 2009), and JMJ14 binding motifs (CTTGnnnnnCAAG) (Zhang *et al*., 2015), no difference was observed between the motifs in the promoters of the genes showing high transcription in WT and in mutants. Additionally, none of the motifs associated with plant hormones, such as cytokinin (ARR10 (GATY)) (Hosoda *et al*., 2002), ethylene (GCC-box (GCCGCC)) (Hao *et al*., 1998), and ABA (ABRE (ACGTG)) (Hattori *et al*., 2002) showed higher occurrence in the promoters of the highly transcribed genes in the *pptrb* mutants.

### Nuclear localization and interaction of TRBs

These findings suggest that PpTRBs in various plant cells (*P. patens*, *A. thaliana*, *N. tabacum* or *N. benthamiana*) predominantly localize to the nucleus. The nuclear localization of PpTRBs in *A. thaliana* protoplasts closely resembles the localization of native AtTRBs in isolated *Arabidopsis* nuclei, as observed using anti-TRB antibodies (Kusová *et al*., 2023). Mutual interactions of PpTRBs were predominantly detected in the nucleus, where they were concentrated within nucleoplasmic foci. This pattern is consistent with the behavior of most AtTRBs in *A. thaliana* and *N. benthamiana* cells (Dvořáčková *et al*., 2010; Kusová *et al*., 2023).

The mutual PpTRB interactions were observed using BiFC, a technique that may also detect indirect interactions, as previously demonstrated for TERT and Recombination UV B-like (RUVBL) proteins, plant homologues of Pontin and Reptin, whose interaction is mediated by TRB proteins (Schořová *et al*., 2019). Additionally, mutual interactions were also confirmed using FLIM-FRET. Since energy transfer between fluorophore groups can only occur efficiently when the interacting proteins are in close physical proximity (≤10 nm), it is highly likely that PpTRBs directly interact. This nuclear localization and interaction pattern supports the hypothesis that TRB proteins have a conserved role in the nucleus, potentially contributing to telomere maintenance and other nuclear processes.

Interestingly, in a subset of *P. patens* protoplasts, mutual PpTRB interactions were also observed in the cytoplasm, particularly near the cellular periphery. This localization pattern resembles that of AtTRB5 in *N. benthamiana*, which is predominantly cytoplasmic (Kusová *et al*., 2023). Furthermore, gene ontology (GO) analysis of moss mutants lacking TRB proteins revealed an enrichment of terms such as ‘cell periphery’ and ‘plasma membrane’. However, the relationship between these phenomena remains unclear and warrants further investigation.

### Evolutionary insights into TRB proteins

A comprehensive phylogenetic analysis encompassing not only mosses, but also Streptophyte algae, hornworts, liverworts, and seed plants demonstrated that genes encoding TRB proteins originated in early-diverging Streptophyta. Transcriptomics data from species lacking assembled genomes were utilized to confirm the absence of detectable TRB gene homologues in the transcriptomes of early-diverging lineages.

In this study, we discovered that Chlorokybaceae, but not Klebsormidiophyceae as we previously mentioned (Kusová *et al*., 2023), is the earlies diverging lineage within Streptophyta in which plant TRB genes could be detected. However, in certain basal representatives, doubts remain regarding the presence of full-length transcripts containing all three defining domains or their correct structure, due to sequence divergence and variability in transcriptome data completeness or quality. Additionally, the reliability of transcriptomic data is influenced by the potential for contamination, which may arise during RNA extraction, sequencing, or deconvolution. To mitigate these issues, we compared transcriptomes from closely related species generated from different Bioprojects, enabling the identification of contamination where possible.

The presence of TRB genes was subsequently newly identified in other streptophyte algal lineages, including Charophyceae, Coleochaetophyceae, and Zygnematophyceae. Among mosses (Bryophyta), the number of TRB genes varied and appeared to be associated with multiple independent ancient WGD events (Gao *et al*., 2022). These events likely occurred across several classes, including Marchantiopsida (liverworts), Sphagnopsida (peat mosses), Bryopsida (joint-toothed mosses), and Anthocerotopsida (hornworts) (Clark and Donoghue, 2018; Gao *et al*., 2022).

In contrast to mosses, seed plants predominantly possess two major diverging lineages of TRBs (Kusová *et al*., 2023). These lineages likely arose from WGD events that expanded the TRB family to as many as ten members, with increased copy numbers observed in polyploid species such as triploids and tetraploids. Unusual branching patterns in seed plants can be explained by the usual phenomena of model inadequacy and pernicious incomplete lineage sorting. Furthermore, these observations could be influenced by errors inherent in automated genome annotation, which remains prone to gene delineation inaccuracies even when trained on transcriptome data (Salzberg, 2019).

Our findings suggest that the evolutionary origins of TRB homologues may predate current estimations. However, a more comprehensive analysis of Chlorophyta and other eukaryotic lineages, such as Excavata, is required. Additionally, genetic manipulation will be necessary to fully elucidate the extent of functional conservation of TRBs across Streptophyta.

## Material and methods

### Accession numbers

PpTRB1 (Pp3c13_1060V3.1); PpTRB2 (Pp3c13_1310V3.1); PpTRB3 (Pp3c3_1630V3.1).

### Plant material cultivation and morphological analysis

The WT ‘Gransden’ strain of *Physcomitrium patens* (Hedw.) B.S.G. (Rensing *et al*., 2008) was used for generation of all *pptrb* mutants. The moss lines were cultured as a ‘spot inocula’ on BCD agar medium supplemented with 1 mM CaCl_2_ and 5 mM ammonium tartrate (BCDAT medium), or as lawns of protonema filaments by subculture of homogenized tissue on BCDAT agar overlaid with cellophane in growth chambers with 18/6 h day/night cycle at 22/18 °C (Cove *et al*., 2009). Homogenized or unhomogenized protonema were grown for one week in normal light conditions on plates where 0.5% sucrose was included in the medium with 1.5% agar and then transferred to the dark for three weeks. Petri dishes were positioned vertically to enhance the observation of caulonema growth.

After 1 month of growth on BCD or BCDAT, whole colonies of WT and *pptrb* mutants, were photographed using by Canon EOS77D camera equipped with a Canon EF 28–135 mm f/3.5–5.6 objective. Gametophore details were captured using a Leica M205FA stereomicroscope with a Plan-Apochromat 2.0x objective. Staining with 10 μg/mL propidium iodide (PI, Sigma-Aldrich) was performed as described previously (Lelkes *et al*., 2023). Images of caulonema growth in the dark were obtained by Epson PERFECTION V700 PHOTO scanner and by Zeiss AxioZoom.V16-Apotome2 stereo zoom microscope with a Plan-Neofluar 1x or a Plan Apo 0.5x objective.

### Construction of *pptrb* mutant lines

The STOP codon knock-in was introduced by homology-directed repair following Cas9-induction of double-strand break within the *PpTRB1* locus. A 50 bp double-stranded DNA donor template containing the desired mutation was used for repair. The donor template was composed of two complementary oligonucleotides (pKA1364 and pKA1365; **Supplementary Table 4**) homologous to the first exon of *PpTRB1* locus. The oligonucleotides introduced a substitution of +63G to C (creating *SalI*I cleavage site), deletion of +69TG, and a substitution of +71G to A, collectively generating a STOP codon at the position of the 23rd amino acid (**Supplementary Figure 1a**). Gateway destination vector pMK-Cas9-gate (Addgene plasmid #113743, which includes a Cas9 expression cassette and kanamycin resistance, along with the entry vector pENTR-PpU6sgRNA-L1L2 (Addgene plasmid #113735) containing the PpU6 promoter, and sgRNA scaffold, were sourced from (Mallett *et al*., 2019). PpTRB1 specific sgRNA spacer was synthesized as two complementary oligonucleotides (pKA1174 and pKA1275**; Supplementary Table 4**). Four nucleotides were added to the 5′-ends of the oligonucleotides to create a sticky ends compatible with *Bsa*I-linearized pENTR-PpU6sgRNA-L1L2 upon annealing. The Cas9/sgRNA expression vector was assembled using the Gateway LR reaction to recombine the entry vector pENTR-PpU6sgRNA-L1L2 with TRB1 sgRNA spacer and destination vector pMK-Cas9-gate. This DNA construct was co-transformed with donor template (annealed pKA1364 + pKA1365) into protoplasts by PEG-mediated transformation (Liu and Vidali, 2011a). The sgRNA was designed in the CRISPOR online software, using *P. patens* (Phytozome V11) as the genome and *S. pyogenes* (5′ NGG 3′) as the PAM parameter. The protospacer with the highest specificity score was selected for further experiments. After five days of regeneration, the transformed protoplasts were transferred to the BCDAT medium supplemented with 50 mg/l G418. Following one week of selection, the G418-resistant lines were propagated. Crude extracts from young tissues of these lines were used for PCR amplification of genomic DNA around editing sites using primers pKA1383, pKA1384. The PCR products were digested by *Sal*I and analyzed by DNA electrophoresis. Lines with cleaved PCR products were subsequently analyzed to confirm the accurate introduction of mutations. Four lines, *pptrb1 #5, #6, #10* and *#12,* were identified with correctly integrated STOP codons. All the lines exhibited very similar growth and morphology phenotypes.

Constructs for gene targeting of *PpTRB2* and *PpTRB3* were assembled in GoldenBraid cloning system (Sarrion-Perdigones *et al*., 2011). Each construct included a selection cassette (NPTII or Hyg^R^) flanked by approximately 1kb of sequences from each of the 5’- and 3’regions of the target genes (**Supplementary Figure 1b, c**). A total of 30μg of linear DNA from each construct was introduced into protoplasts using PEG-mediated transformation (Liu and Vidali, 2011a). After five days of regeneration, the transformed protoplasts were transferred to Petri dishes containing the BCDAT medium supplemented with 30 mg/l hygromycin or 50 mg/l G418. Following three rounds of selection, the transformants were deemed stable. Gene replacement was confirmed by PCR of genomic DNA isolated from the stable transformants, following the method described by (Dellaporta *et al*., 1983). Primers were designed to amplify the selection cassette in an outward direction, pairing with gene-specific primers that annealed to sequences outside the homology regions integrated into the targeting vectors (**Supplementary Figure 1b, c; Supplementary Table 4**). A single representative line for each confirmed mutant, *pptrb1 #5*, *pptrb2 #11 and pptrb3 #2*, was selected for further investigation.

Double mutants were prepared and verified as described above, as follows: (i) *pptrb1 pptrb2* mutants were prepared by delivering of *PpTRB1* CRISPR/Cas9 construct into protoplasts isolated from *pptrb2 #11*; (ii) *pptrb1 pptrb3* mutants were prepared by delivering of *PpTRB1* CRISPR/Cas9 construct into protoplasts isolated from *pptrb3 #2*; (iii) *pptrb2 pptrb3* mutants were prepared by delivering of *PpTRB3* replacement vector into protoplasts isolated from *pptrb2 #11*. From two to three independent double mutant lines were investigated, and a single representative line was subsequently selected for further detailed investigation: *pptrb1 pptrb2 #3*, *pptrb1 pptrb3 #6* and *pptrb2 pptrb3 #16*.

### RNA isolation and QuantSeq 3’mRNA-seq

The total RNA was isolated from 7-day-old protonemal tissue (WT, *pptrb1 #5, pptrb2 #11, pptrb3 #2, pptrb1 pptrb2 #3, pptrb1 pptrb3 #6, pptrb2 pptrb3 #16*). Approximately 100 mg of tissue per sample was homogenized using glass beads. 1 ml of TRI reagent (TRI REAGENT®; MRC) was added and incubated 5 min at room temperature. Then, 200 µl of chloroform was added and mixed, followed by centrifugation for 15 min at 12 000 g at 4°C. After centrifugation, the upper aqueous phase (appr. 500 µl) was used for RNA purification with Direct-zol RNA Miniprep Kits (Zymo Reasearch) including DNaseI treatment. The integrity of RNA was checked by agarose gel electrophoresis, and concentration was measured using a NanoDrop spectrophotometer (NanoDrop Technologies).

RNAseq analysis was carried out by Core Facility Genomics of CEITEC Masaryk University. RNA integrity was checked on the Fragment Analyzer using RNA Kit 15 nt (Agilent Technologies). 400 ng of total RNA was used as an input for library preparation using QuantSeq 3’mRNA-Seq Library Prep Kit (FWD) (Lexogen) with polyA selection in combination with UMI Second Strand Synthesis Module for QuantSeq FWD and Lexogen i5 6 nt Unique Dual Indexing Add-on Kit (Lexogen). Library quantity and size distribution was checked using QuantiFluor dsDNA System (Promega) and High Sensitivity NGS Fragment Analysis Kit (Agilent Technologies). Final library pool was sequenced on NextSeq 500 (Illumina) using High Output Kit v2.5 75 Cycles (Illumina), resulting in an average of 12 million reads per sample.

### Data analysis and gene onthology

The Binary Base Call (BCL) files obtained from QuantSeq 3’mRNA-seq were converted to FASTQ format using *bcl2fastq* v. 2.20.0.422 Illumina software for base calling. Quality check of raw single-end FASTQ reads was carried out by FastQC (1). The adapters and quality trimming of raw FASTQ reads was performed using Cutadapt v4.3 (Martin, 2011) with Illumina adapter trimming and parameters -m 35 -q 0,20. Trimmed RNA-Seq reads were mapped against the genome of *Physcomitrium patens* (Ppatens_870_v6.1) using STAR v2.7.11 (Dobin *et al*., 2013) as splice-aware short read aligner and default parameters except --outFilterMismatchNoverLmax 0.4 and --twopassMode Basic.

The differential gene expression analysis was calculated based on the gene counts produced using featureCounts from Subread package v2.0 (Liao *et al*., 2014) and further analyzed by Bioconductor package DESeq2 v1.34.0 (Love *et al*., 2014). Data generated by DESeq2 with independent filtering were selected for the differential gene expression analysis due to its conservative features and to avoid potential false positive results, principal component analysis (PCA) was computed from gene expression normalized using DESeq2 Variance Stabilizing Transformation (VST). Genes were considered as differentially expressed (DEG) based on a cut-off of adjusted p-value < 0.05 and log2(fold-change) ≥1 or ≤-1. TPM values were computed using DGEobj.utils convertCounts function. Clustered heatmaps were generated from selected top differentially regulated genes using R package pheatmap v1.0.12, volcano plots were produced using ggplot2 v3.3.5 package and MA plots were generated using ggpubr v0.4.0 package.

Gene Ontology (GO) enrichment analysis was done by g:Profiler version e109_eg56_p17_1d3191d (Raudvere *et al*., 2019, https://biit.cs.ut.ee/gprofiler) with the default setting using DEGs as input, and the number of terms used for plotting was reduced based on the results from REVIGO (Supek *et al*., 2011, http://revigo.irb.hr/) with the predefined cutoff value C = 0.5.

### Cloning of constructs encoding PpTRB proteins

To obtain amplicons with *PpTRB1, PpTRB2 and PpTRB3* genes, total RNA was isolated from 7-day-old protonemal tissue using TRI reagent (TRI REAGENT®; MRC). cDNA was synthetized from 1 µg of RNA via reverse transcription using SMARTScribe™ Reverse Transcriptase (TAKARA) and oligo dT-VN primer (Clontech) (**Supplementary Table 4)**. Expression vectors were constructed using the Gateway® system (Invitrogen). The constructs encoding PpTRB proteins were amplified from cDNA using KAPA Taq DNA Polymerase (KAPA Biosystems) and specific primers F-PpTRB1 and R-PpTRB2(stop) for PpTRB1, F-PpTRB2 and R-PpTRB2(stop) for PpTRB2, and F-PpTRB3 and R-PpTRB3(stop) for PpTRB3 (**Supplementary Table 4**). PCR products were adapted for Gateway cloning in adaptor PCR using attB1/2 primers (Invitrogen) and KAPA Taq DNA Polymerase, and introduced directly to entry Gateway vectors (pDONRZeo, pDONR207) using the BP Clonase™ II enzyme mix (Thermo Fisher Scientific). Alternatively, PCR products were introduced into pCR™II-TOPO® vector (TOPO® TA Cloning® Kit (Invitrogen)) and sequenced. TOPO plasmids then served as templates in adaptor PCR or their inserts were recombined to entry vectors using the BP Clonase™ II enzyme mix (Thermo Fisher Scientific) according to manufacturer’s instruction. To generate expression clones, all entry clones were used in LR Clonase™ II (Thermo Fisher Scientific) reactions with corresponding expression vectors: pSAT5-DEST-c(175-end)EYFP-C1(B) and pSAT4-DEST-n(174)EYFP-Cl for BiFC assay (Lee *et al*., 2012), pH7WGF2 and pB7WGR2 for fusion with GFP/RFP, respectively (Karimi *et al*., 2002).

### Plant cells transfection

For protein localization, expression clones of *PpTRB* genes in pH7WGF2 vector containing eGFP protein coding sequence were used (Karimi *et al*., 2002). The localization of PpTRB proteins was analyzed in *P. patens* protonema protoplasts, *A. thaliana* root protoplasts and *N. tabacum* leaves.

*P. patens* protoplasts were prepared and transfected using PEG-based procedure as described in (Liu and Vidali, 2011a) from 7-day-old protonemata grown on BCDAT agar medium. For GFP localization and BiFC analysis (see below), 10 µg of plasmid DNA for each construct was used.

*A. thaliana* root protoplasts were isolated and transfected with 10 µg of plasmid DNA per construct using PEG-mediated method as described in (Kolářová *et al*., 2021).

Transient heterologous expression in *N. tabacum* (SR1 Petit Havana) leaves was performed as described in (Kusová *et al*., 2023). The leave epidermal cells were transfected by the infiltration procedures described in (Voinnet *et al*., 2000).

The fluorescence was observed using a laser scanning confocal microscope Zeiss LSM 880 with C-Apochromat 63x/1.2 W Korr M27 objective using 488 nm laser for GFP.

### Protein localization and FLIM-FRET

For FLIM-FRET analysis, coding sequences of PpTRB proteins fused to GFP/RFP (pH7WGF2/pB7WGR2) were used. *A. tumefaciens* competent cells (strain GV3101) were transfected with corresponding expression clones and these plasmids were used for transient expression in *N. tabacum* (SR1 Petit Havana) leaves (Voinnet *et al*., 2000). Laser scanning confocal imaging microscope Zeiss LSM 780 AxioObserver equipped with external In Tune laser (488-640 nm, < 3nm width, pulsed at 40 MHz, 1.5 mW) C-Apochromat 63 x water objective, NA 1.2 was employed for confocal microscopy. HPM-100-40 Hybrid Detector from Becker and Hickl GmbH including Simple-Tau 150N (Compact TCSPC system based on SPC-150N) with DCC-100 detector controller for photon counting was used for FLIM-FRET data acquisition. Zen 2.3 light version from Zeiss was used for processing confocal images. The acquisition and analysis of FLIM data involved the utilization of SPCM 64 version 9.8. and SPCImage version 7.3 from Becker and Hickl GmbH, respectively. A multiexponential decay model was used for fitting.

### Bimolecular Fluorescence Complementation (BiFC)

To analyze protein-protein interactions by BiFC, the *P. patens* protoplasts were prepared and transformed with expression clones of *PpTRB* genes in pSAT4-DEST-n(174)EYFP−Cl (pE3136/nYFP) and pSAT5-DEST-c(175-end)EYFP−C1(B) (pE3130/cYFP) vectors. To control PEG-based transfection efficiency and to label cell nuclei, the protoplasts were co-transfected with a plasmid expressing mRFP fused to the nuclear localization signal of the VirD2 protein (mRFP−VirD2NLS) (Citovsky *et al*., 2006). The transformation was performed as described by (Liu and Vidali, 2011b) with minor modifications. After the incubation period of protoplasts with PEG solution, the protoplasts were diluted with 3 mL of MMg solution instead of W5 solution, then centrifuged at 250 g for 5 minutes and finally resuspended in 600 µl of MMg solution. Transfected protoplasts were incubated in the light, at room temperature overnight. The next day, the fluorescence was observed using a laser scanning confocal microscope Zeiss LSM 880 with C-Apochromat 63x/1.2 W Korr M27 objective using 488 nm laser for GFP, 514 nm laser for YFP and 594 nm laser for RFP. The images obtained were adjusted in the ZEN 2.5 lite Blue software.

### Terminal Restriction Fragment Analysis (TRF)

Genomic DNA (gDNA) from WT and mutant lines was isolated from 7-day-old protonemal culture (WT, *pptrb1 #5, pptrb2 #11, pptrb3 #2, pptrb1 pptrb2 #3, pptrb1 pptrb3 #6, pptrb2 pptrb3 #16*) according to (Dellaporta *et al*., 1983). The quality of DNA was checked, and its concentration determined by electrophoresis in a 1% (w/v) agarose gel stained with ethidium bromide using Gene Ruler 1 kb DNA Ladder (Thermo Scientific, Waltham, MA) as standard and Multi Gauge software (Fujifilm, Tokyo, Japan).

Telomere lengths were analysed as described in (Fojtová *et al*., 2015). Samples of 1 µg genomic DNA were digested by 1 µl of MseI overnight (10,000 units/ml, NEB). The next day, an additional 1 µl of MseI was added to samples and the digestion continued for about 4 hours. Then, samples were separated in 1% (w/v) agarose gel followed by Southern blot on Amersham™ Hybond™ - XL membrane (Ge Healthcare, Chicago). The membrane was hybridized overnight at 65°C (to avoid signals from interstitial telomeric sequences) with plant telomere probe. The probe was synthesized by non-template PCR and radioactively labelled with [^32^P-dATP] using DecaLabel DNA labelling kit (Thermo Scientific). Signals were visualized using FLA7000 imager (Fujifilm). Boxplot values were obtained by evaluating telomere length profiles of mutant and corresponding wild-type plants using R-based online toolset WALTER (Lyčka *et al*., 2021), using default setup with background correction.

### Phylogenetic analysis

To effectively identify TRB orthologs across the Streptophyta clade, we deployed an iterative approach, wherein after each step, the query dataset was enriched with the newly discovered protein sequences. The resulting multiple sequence alignment (MSA) was then used to target lineages lacking positive hits. The iteration proceeded until all relevant data had been processed.

Previously discovered TRB sequences (Kusová *et al*., 2023) were queried against the NCBI nr database (MMseqs Version: 14.7e284) and restricting output to alignments that are within the 1.0E-06 threshold e-value with default sensitivity.

The retrieved Streptophyta protein sequences were aligned with FAMSA 2 (Deorowicz *et al*., 2016) and converted into a MMseqs2 profile. Relying on iterative profile-to-sequence search in MMseqs2, we probed additional taxa not included in nr. Taxa were selected to cover missing lineages in Streptophyta. In case of annotated genome, we used translated gene sequences (proteome), otherwise we selected taxa with available transcriptome data. We used publicly available RNAseq data for 30 species. To remove poor quality reads, polyAtails, and singletons, reads were filtered with fastp (Chen *et al*., 2018; Chen, 2023) running on the following settings: --trim_poly_g --trim_poly_x --low_complexity_filter -- cut_tail. The filtered read sets were assembled using rnaSPAdes v4.0.0 with default options (Bushmanova *et al*., 2019). De novo assembled transcriptomes and previous assemblies from 1KP (One Thousand Plant Transcriptomes Initiative, (Leebens-Mack *et al*., 2019)) were searched with MMseqs2. When medium sensitivity screening failed to retrieve any bona fide matches, we repeated the search in high sensitivity mode (-s 7.5). Protein-coding open reading frames were identified from the profile alignments. For annotated genomes lacking protein sequences, we obtained the proteome from CDS extraction with gffread (Pertea and Pertea, 2020) and translation with orfipy release 0.0.4 (Singh and Wurtele, 2021) keeping the longest open reading frames (ORFs).

When no positive hits were recovered from transcriptome nor proteome data, we turned our attention to readily available unannotated genomes. We defined TRB gene candidate regions with two complementary approaches: 1) using miniprot (Li, 2023) for mapping previously identified proteins to the genomes and improved exon delimitation with Miniprot boundary scorer (https://github.com/tomasbruna/miniprot-boundary-scorer) (Brůna *et al*., 2023), 2) the MSA from the recovered proteins was used to build a profile hidden Markov models for BATH (Krause *et al*., 2024) and genomes were probed with the bathsearch module. We retained genes that were validated by the highest scoring predictions from both approaches and translated them with orfipy. Transcriptomes, proteomes and genomes were assessed for completeness by running BUSCO v 5.8.0 in the appropriate mode against the Viridiplantae databases (viridiplantae_odb10) (Seppey *et al*., 2019). All newly identified protein sequences were checked against the InterPro 103.0 database (Apweiler *et al*., 2001) for the presence of Myb-like, H1/5-like and coiled-coil domains in a N to C-terminal orientation.

As the broad evolutionary patterns of TRB evolution have been previously studied in Angiospermae (Kusová *et al*., 2023), the MSA was downsized for phylogenetic purposes to a single taxon per family in this clade. We kept taxa with the highest TRB copy number and retained all *Brassicaceae* sequences in order to get a deeper insight into the gene diversification rate within *Arabidopsis thaliana* relatives.

ModelFinder (Kalyaanamoorthy *et al*., 2017) was used to determine the best fitting molecular evolutionary models for maximum likelihood analyses. Protein mixture and nonreversible models (Minh *et al*., 2021) were tested alongside the base time-reversible amino acid models. Selected models were used for building the phylogenies in the IQ-TREE software package (Nguyen *et al*., 2015); branch supports was assessed with ultrafast bootstrap approximation with 10000 replicates (Hoang *et al*., 2018). ModelFinder recovered that the reversible JTT+I+R8 protein evolution model was a better fit than the LG+C10+F+I+G mixture model that was ranked close second according to the Bayesian Information Criterion (164429.949 vs 164435.425). We attempted to infer the most likely root placement using the non-reversible unrestricted model (UNREST) in IQTREE. Because of low rootstrap support values (21.4%), the root location was unresolved and consequently we rooted the tree with the Chlorokybophyceae lineage as outgroup, in accordance with the accepted phylogeny of Streptophyta. Since outgroup rooting might distort ingroup topology due to long-branch attraction where distantly related or fast-evolving taxa spuriously branch with the outgroup (Philippe *et al*., 2011), we compared the topology with the tree obtained from a reduced dataset without outgroup sequences. The two typologies were perfectly congruent, thus dismissing a reconstruction error arising from distant outgroup inclusion. Trees were visualized in iTOL v6 (Letunic and Bork, 2024).

## Author contribution

P.P.S. and J.F. conceived the study. E.S. performed the cloning, and A.Ku. performed localization and protein-protein interaction studies. Telomere measurements were performed by A.Ku. with the help of I.G.P., and RNA-seq analysis was carried out by A.Ku. with the help of J.R. FLIM-FRET was performed by J.S. with the help of A.Ku. and J.H. TPM calculations for GOLEM were done by B.K., with the help of D.H. T.P. and A.Ku. helped with GOLEM program analysis. The mutant plants were established by M.H., with the help of D.Z. The evolution study performed Y.J.K.B.. P.P.S. wrote the paper with help of all co-authors.

## Acknowledgments

Plant Sciences Core Facility of CEITEC Masaryk University is acknowledged for technical support. We acknowledge the core facility CELLIM supported by MEYS CR (LM2023050 Czech-BioImaging). We also acknowledge CF Genomics and CF Bioinformatics supported by the NCMG research infrastructure (LM2023067 funded by MEYS CR) for their support with obtaining scientific data presented in this paper. We acknowledge the Imaging Facility of the Institute of Experimental Botany AS CR supported by the MEYS CR (LM2023050 Czech-BioImaging) and IEB AS CR. For evolutionary analyses, the computational resources were provided by the e-INFRA CZ project (ID:90254), supported by the Ministry of Education, Youth and Sports of the Czech Republic.

