## Supplementary Figures for "Molecular and Phenotypic Characterization of Telomere Repeat Binding (TRBs) Proteins in Moss: Evolutionary and Functional Perspectives"

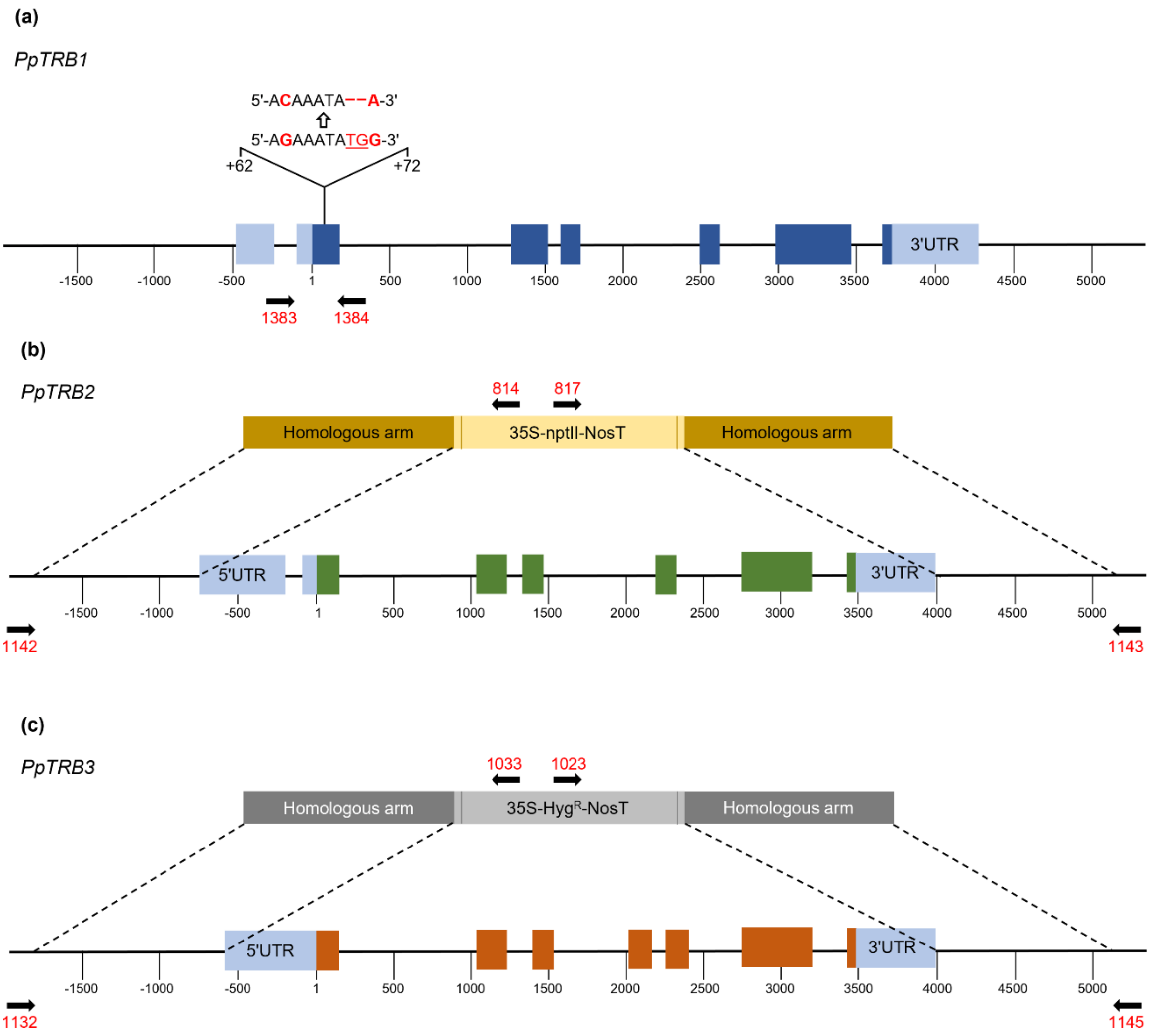


**Supplementary Figure 1. Schematic representation of mutant lines. (a)** Diagram of *PpTRB1* locus indicating the site of incorporated mutations. Nucleotide positions are given relative to the ATG codon, with adenine designated as position +1. Diagrams of *PpTRB2* **(b)** and *PpTRB3* **(c)** loci showing the locations of knockout cassettes containing homologous arms flanking the selectable marker gene, along with positions of genotyping primers. Notably, non-specific transcriptional upregulation downstream of the 35S-HygR-NosT selection cassette — used for homologous recombination-based mutagenesis — was observed in *pptrb2* and *pptrb3* lines.

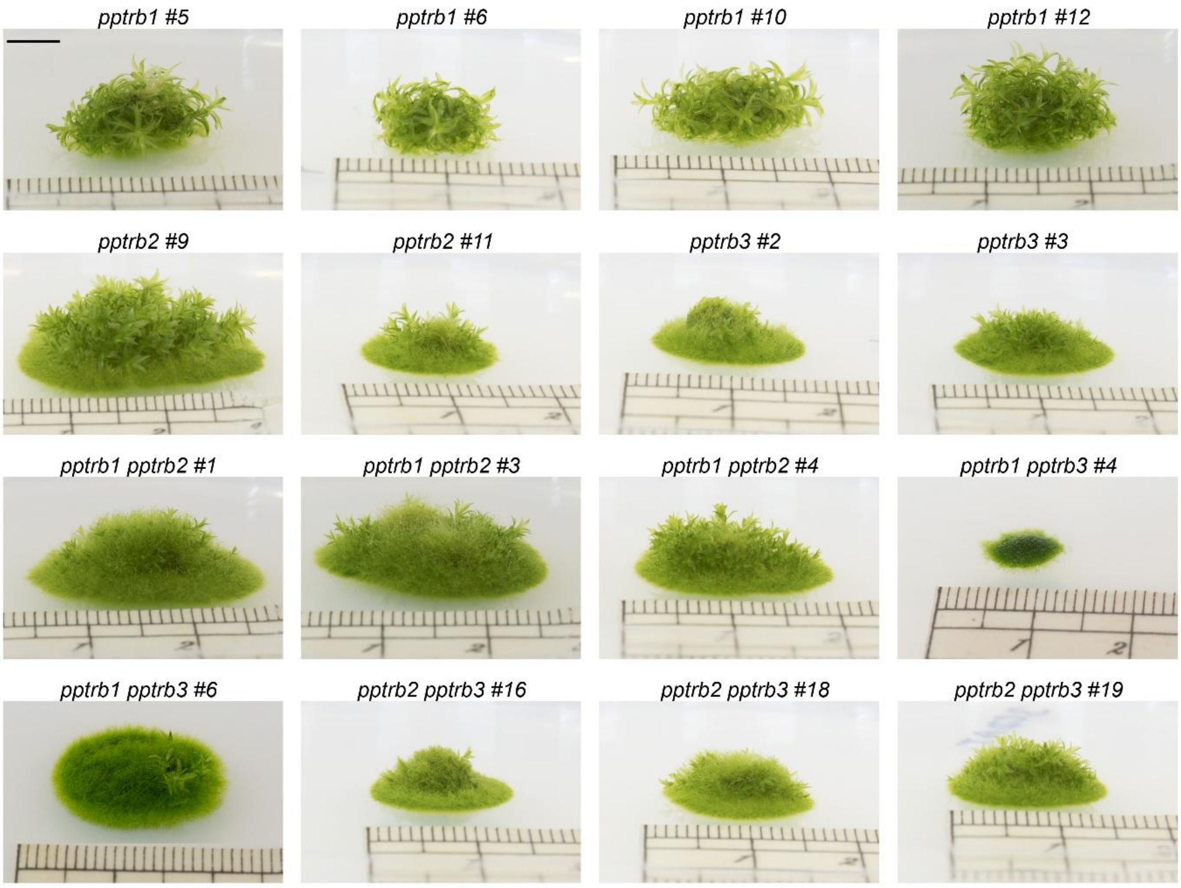


**Supplementary Figure 2. Representative images of all verified mutant lines, prepared as described in the Methods section.** For the mutant analysis, multiple lines were generated for each representative mutant (single and double). Single representative line was selected for further investigation and is also depicted in Figure 1: *pptrb1 #5*, *pptrb2 #11*, *pptrb3 #2*, *pptrb1 pptrb2 #3*, *pptrb1 pptrb3 #6* and *pptrb2 pptrb3 #16*.

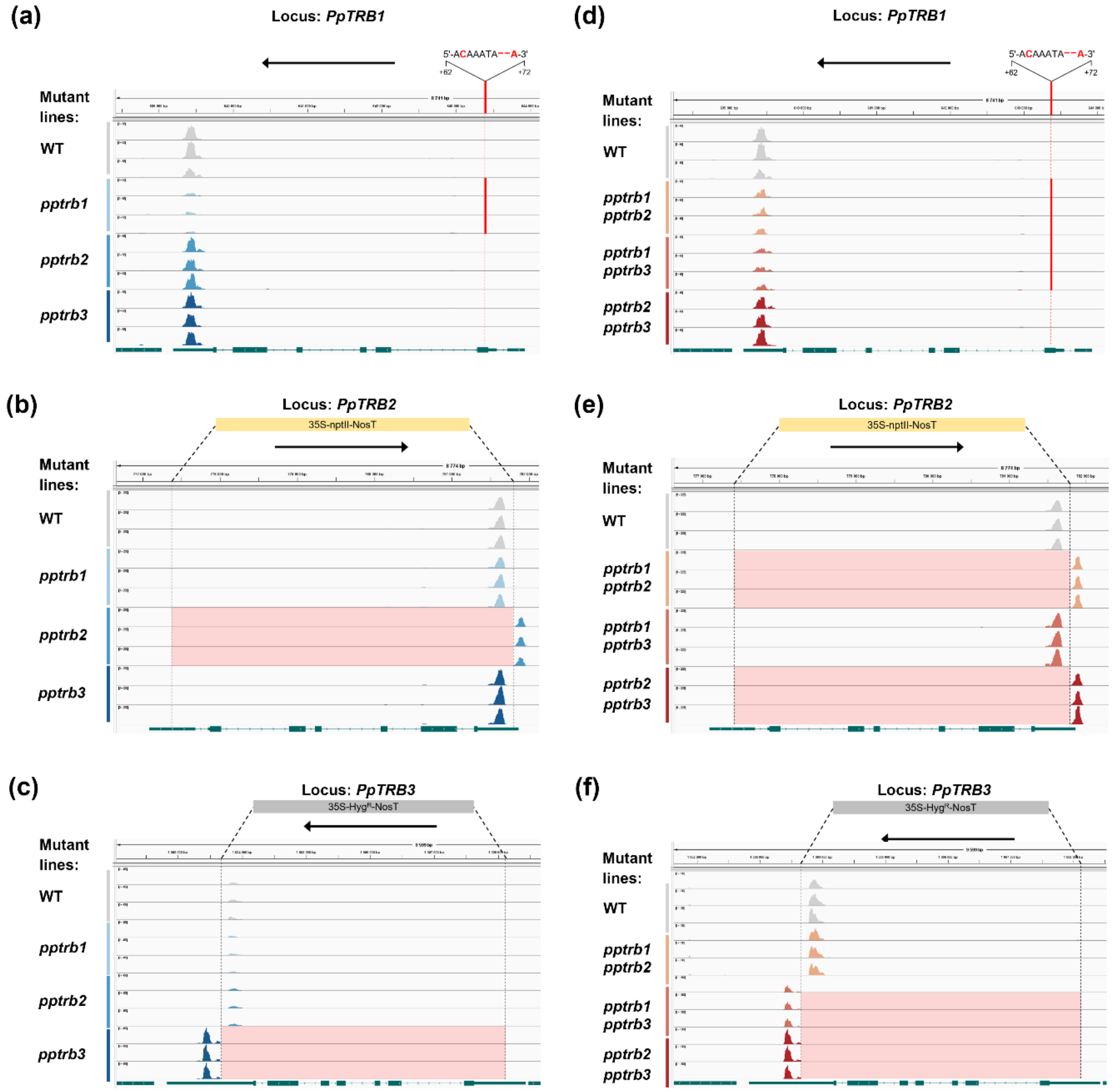


**Supplementary Figure 3. validation of disrupted gene transcription of in *pptrb* lines using QuantSeq.** RNA-Seq QuantSeq, which selectively captures the 3' ends of polyadenylated transcripts, was employed to assess transcript abundance in *PpTRB1*, *PpTRB2*, and *PpTRB3* loci. Sequencing reads were aligned to the corresponding genomic regions, and the results were visualized using the Integrative Genomics Viewer (IGV). Schematic diagrams of the mutations and deletions characterizing each *pptrb* mutant line are presented alongside the mapped read profiles. IGV peak patterns revealed a clear knock-down of *PpTRB1* expression in *pptrb1* mutants and complete knock-out of *PpTRB2* and *PpTRB3* expression in *pptrb2* and *pptrb3* mutant lines, respectively.

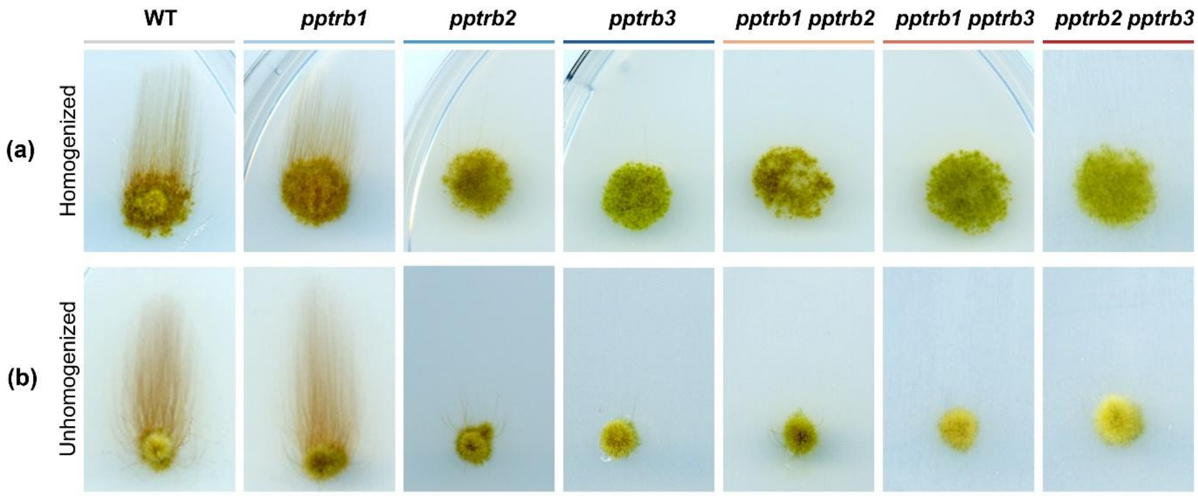


**Supplementary Figure 4. The transition of chloronema to caulonema in the dark.** Mosses grown under light conditions for one week were transferred to darkness for three weeks. Under these conditions, WT plants developed long, negatively gravitropic caulonemata, while *pptrb2*, *pptrb3*, and all double mutants exhibited a significant reduction in caulonemal growth. **(a)** A 50 µl aliquot of the homogenized protonema, collected during the mosses' passaging, was spotted onto the plate. **(b)** A small piece (approximately 1 $mm^2$) from a one-week-old colony was transferred onto the plate.

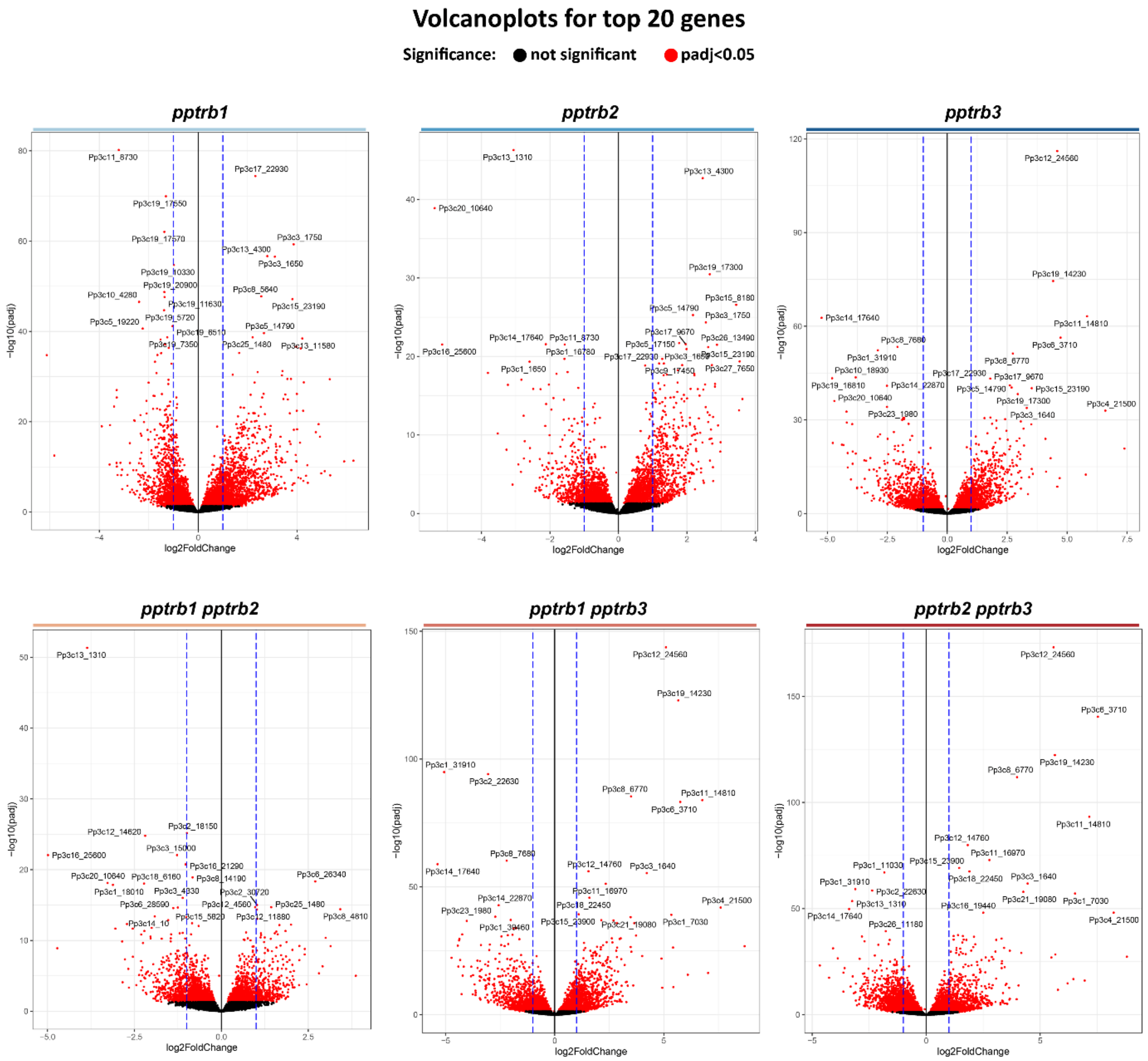


**Supplementary Figure 5. Volcano plots of differentially expressed genes between WT and *pptrb* single and double mutant lines.** Volcano plots illustrate the differential gene expression profiles between wild-type (WT) and *pptrb* mutant lines. The top 20 differentially expressed genes are labeled with their corresponding *Pp* gene identifiers. Genes were considered significantly differentially expressed if they met the thresholds of $p < 0.05$ and absolute fold change > 1. Red dots represent significantly altered genes, while black dots denote genes that do not meet the significance criteria.

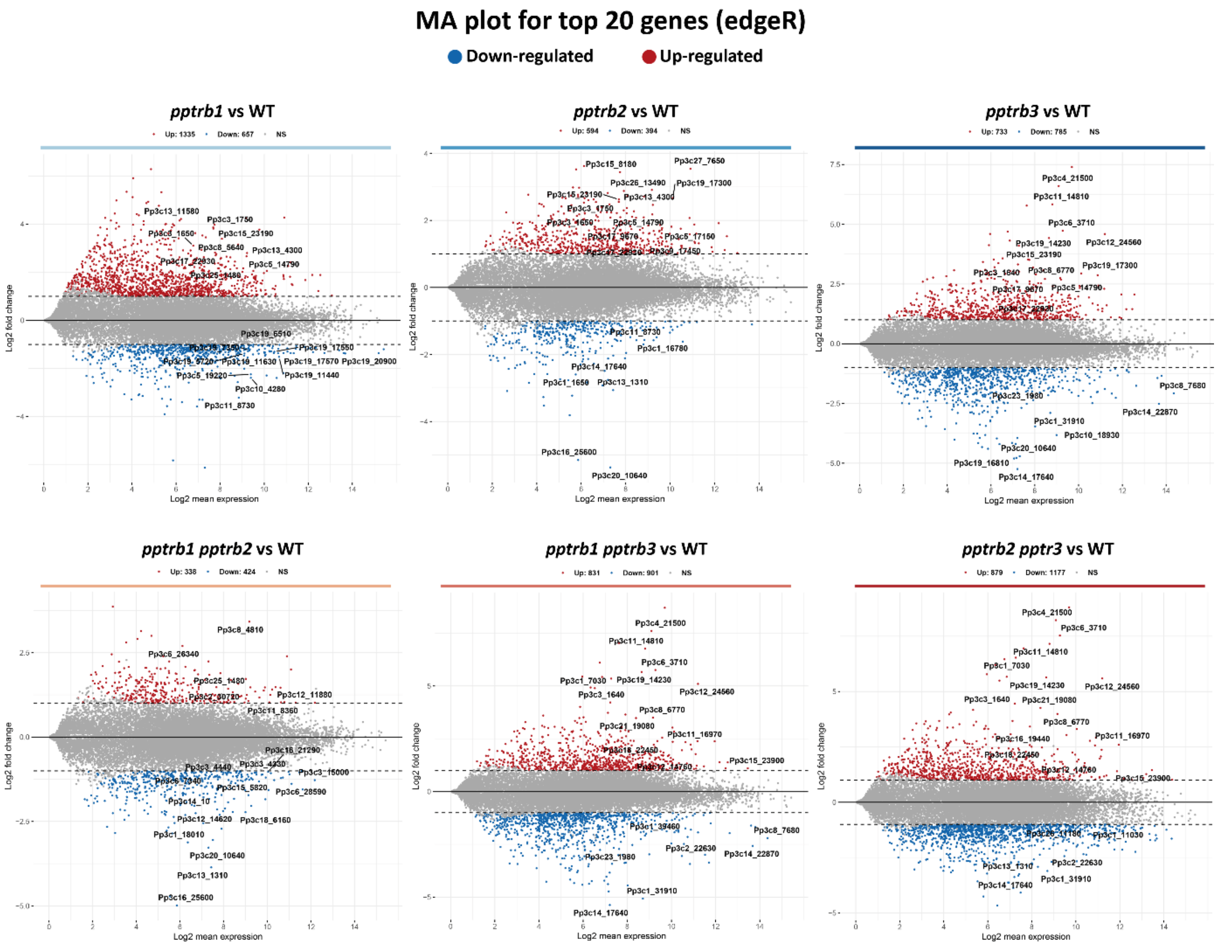


**Supplementary Figure 6. MA plots of differentially expressed genes between WT and *pptrb* single and double mutant lines.** MA plots display the $log_2$ fold changes in gene expression (y-axis) plotted against the mean expression levels (x-axis) for comparisons between wild-type (WT) and *pptrb* mutant lines. The top 20 differentially expressed genes are highlighted and labeled with their respective *Pp* gene identifiers.

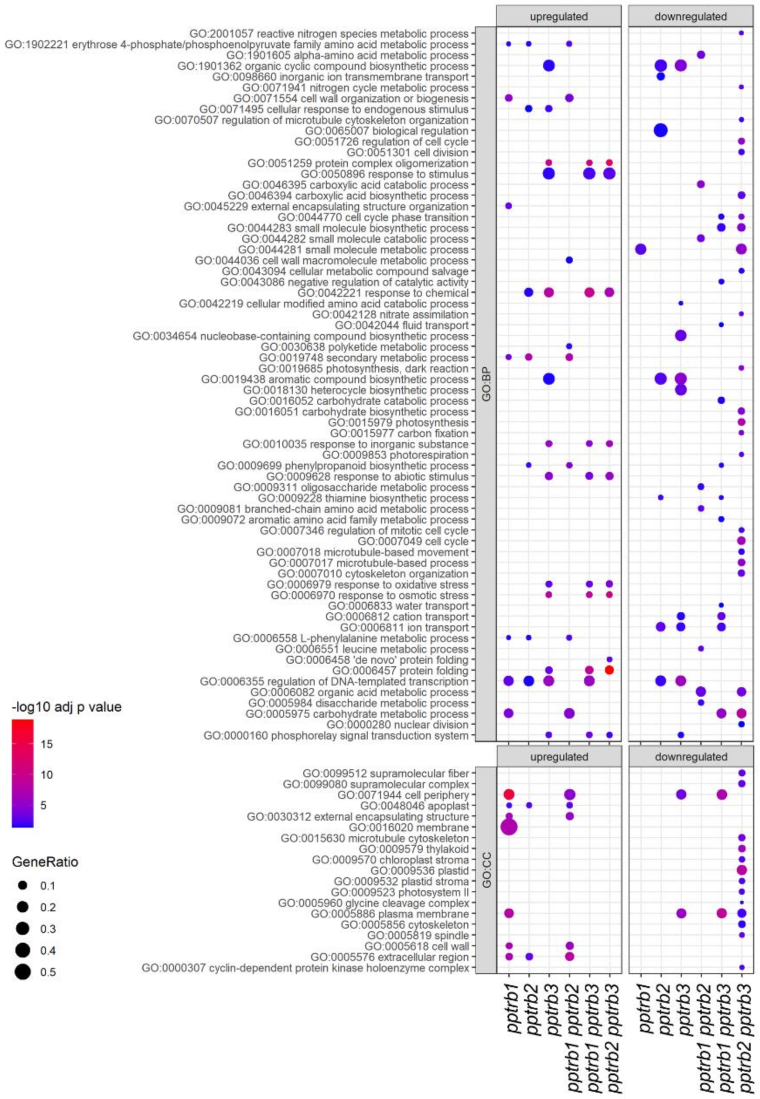


**Supplementary Figure 7. Gene Ontology (GO) enrichment analysis of differentially expressed genes in *pptrb* mutant lines.** GO enrichment analysis was performed on differentially expressed genes (DEGs) using g:Profiler. To reduce redundancy, the resulting GO terms were filtered using REVIGO with a similarity cutoff value of *C* = 0.5. Adjusted *p*-values for each GO term were obtained from g:Profiler. The GeneRatio represents the proportion of DEGs annotated to a given GO term and was calculated as the ratio of *intersection_size* to *query_size*, as defined by g:Profiler.

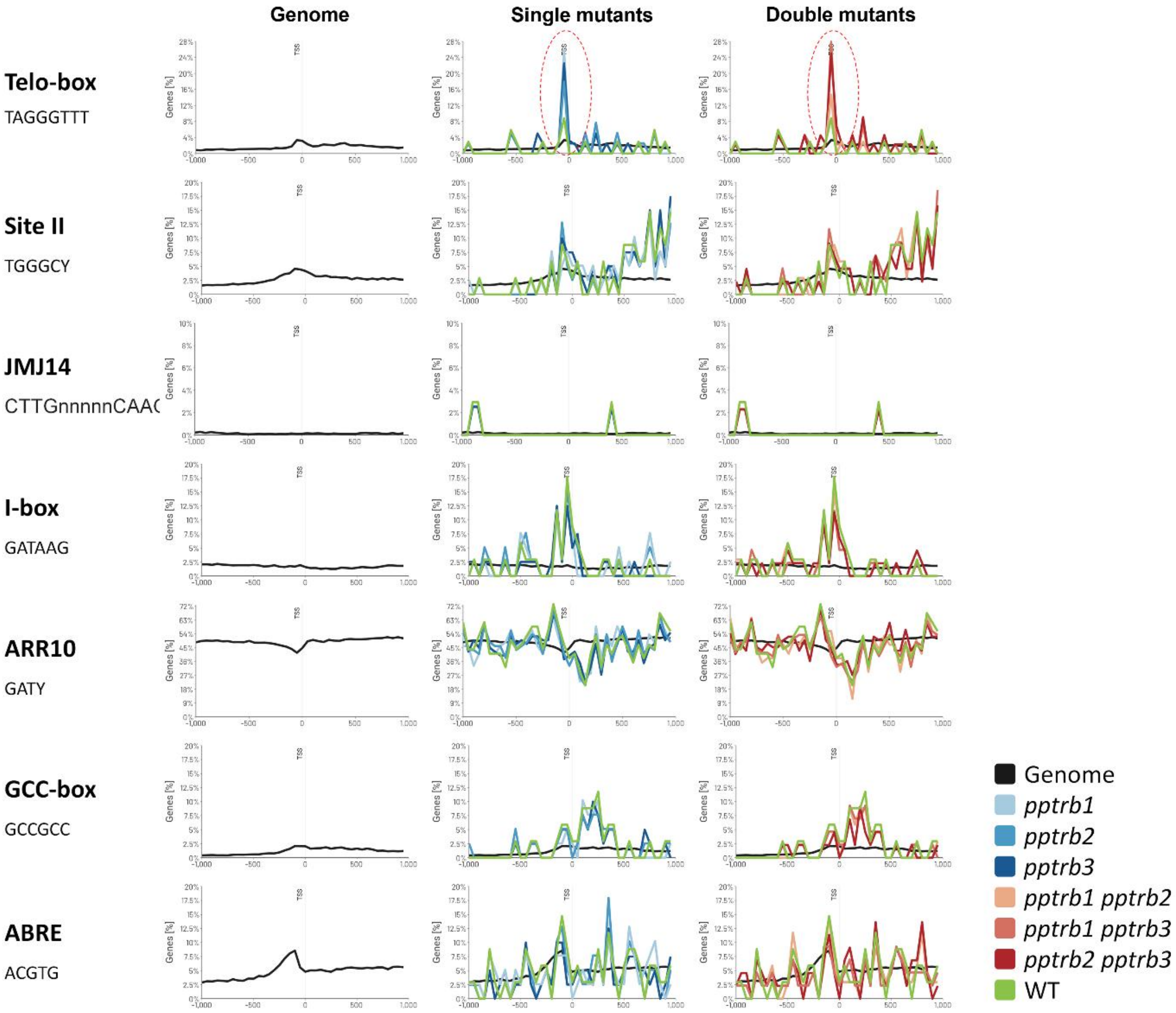


**Supplementary Figure 8. Analysis of RNAseq data from single and double *pptrb* mutant lines in GOLEM web-tool for the presence of *cis*-regulatory motifs.** The GOLEM tool (Nevosád et al., 2025) was used to analyze RNA-Seq data from wild-type (WT) and *pptrb* mutant plants for the presence and positional distribution of cis-regulatory motifs relative to the transcription start site (TSS). The analysis revealed several regulatory motifs in the promoter regions of genes with the highest expression levels in *pptrb* mutants. Notably, the *telo*-box motif was more frequently observed in promoters of highly expressed genes in *pptrb* mutants compared to WT. In contrast, the JMJ14 and Site II motifs—previously linked to TRB protein function—did not show enrichment in *pptrb* mutants.

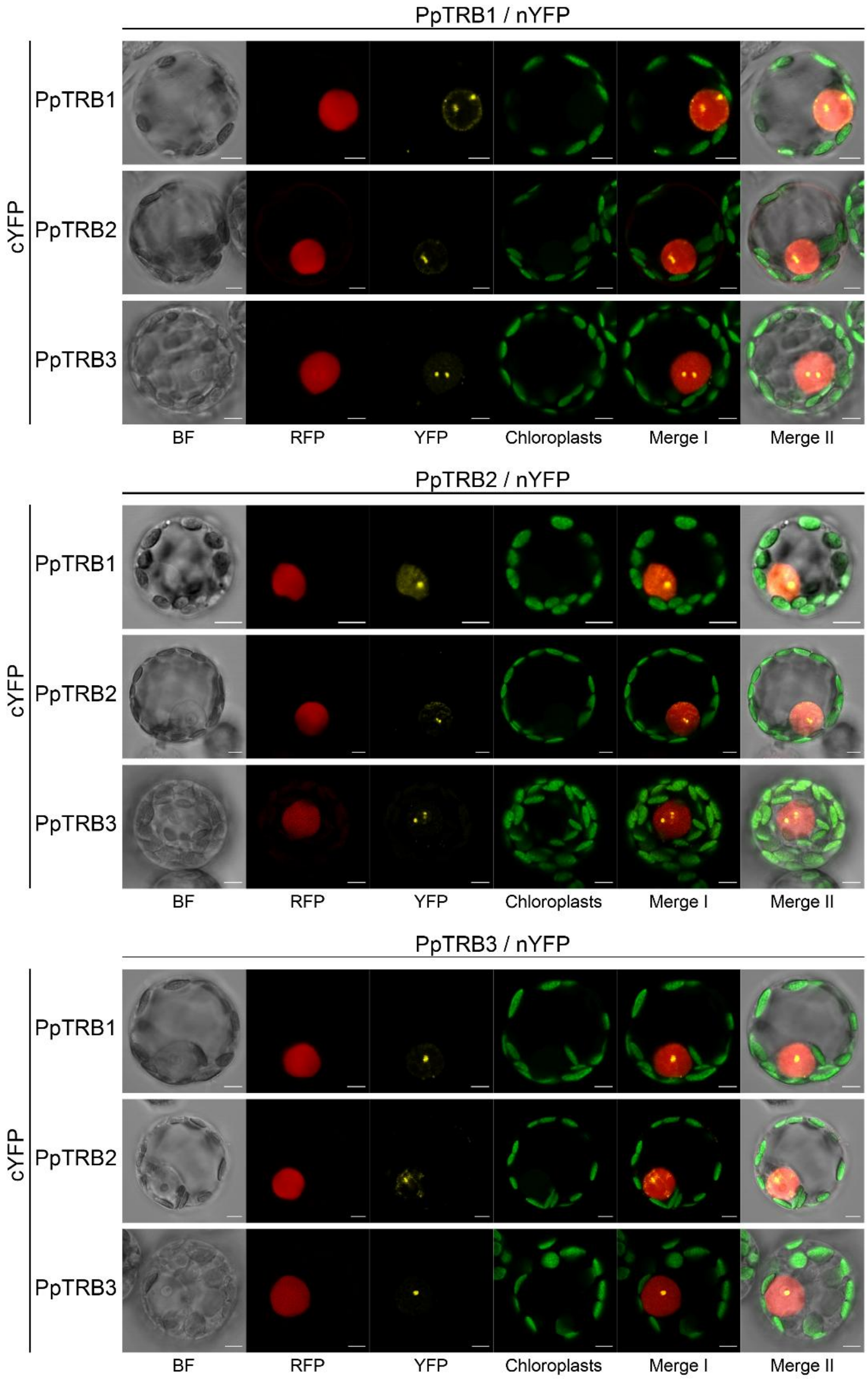


**Supplementary Figure 9. PpTRBs mutually interact in the nucleus.** Protein–protein interactions among *PpTRB* proteins were analysed using a Bimolecular Fluorescence Complementation (BiFC) assay. *PpTRB* constructs fused to either the N-terminal (nYFP) or C-terminal (cYFP) fragments of YFP were co-expressed in protoplasts derived from 7-day-old *P. patens* protonema. Fluorescence signals were visualized in individual channels. YFP channel images reveal distinct fluorescent foci localized within the nucleus, indicating physical interactions between *PpTRB* proteins. *Merge I* displays a merged channels of RFP, YFP, and chloroplast autofluorescence. *Merge II* includes bright-field (BF) in addition to RFP, YFP, and chloroplast autofluorescence channels. Scale bars = 5 µm.

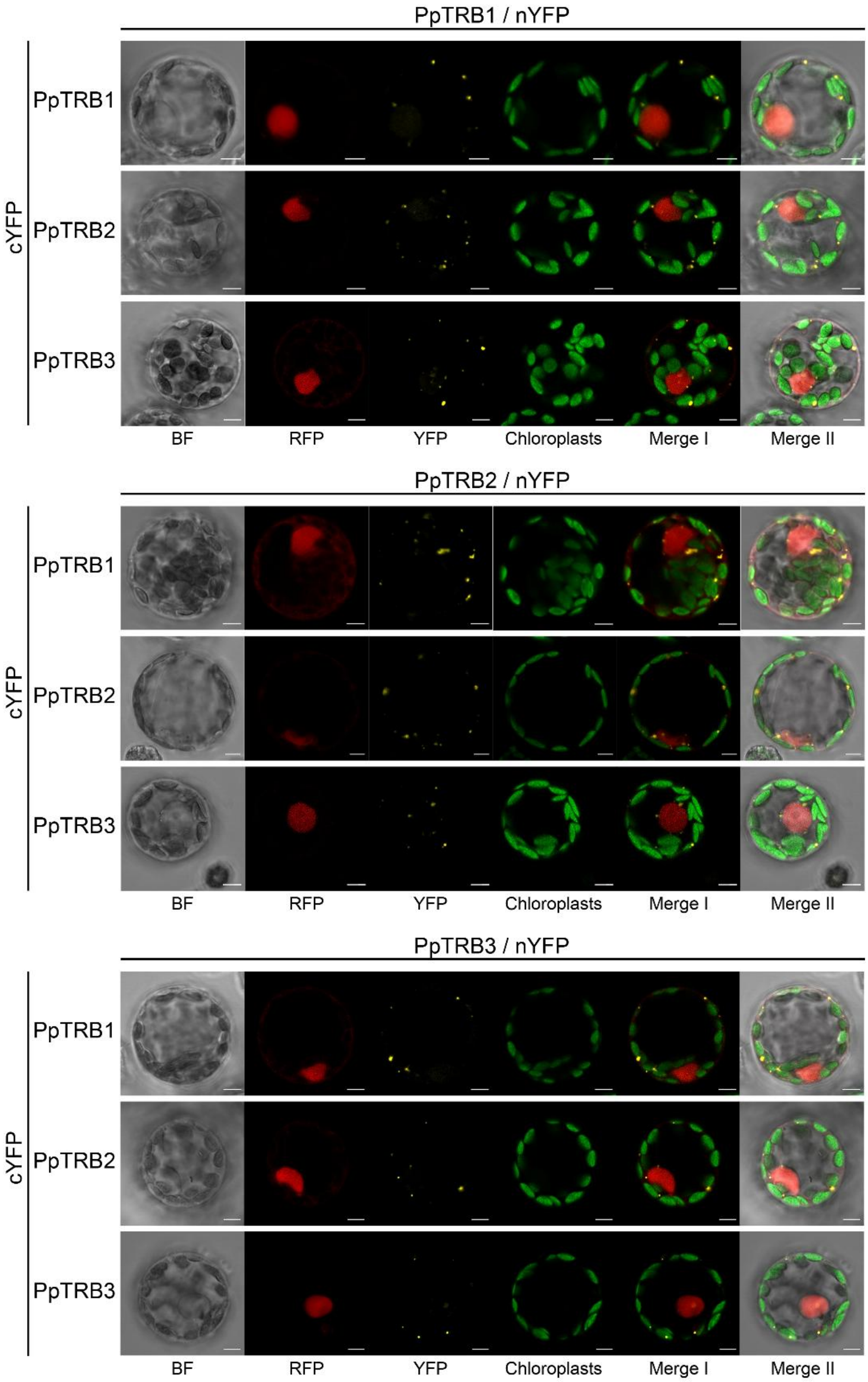


**Supplementary Figure 10. PpTRBs mutually interact in the cytoplasm.** Protein–protein interactions among *PpTRB* proteins were investigated using a Bimolecular Fluorescence Complementation (BiFC) assay. Constructs encoding *PpTRBs* fused to either the N-terminal (nYFP) or C-terminal (cYFP) fragments of YFP were co-expressed in protoplasts isolated from 7-day-old *P. patens* protonema. Fluorescence signals are shown as individual channels. In the YFP channel, fluorescent foci are observed in the cytoplasm, indicating interactions occurring outside the nucleus. *Merge I* shows the overlay of RFP, YFP, and chloroplast autofluorescence signals. *Merge II* includes bright-field (BF) along with RFP, YFP, and chloroplast autofluorescence. Scale bars = 5 µm.

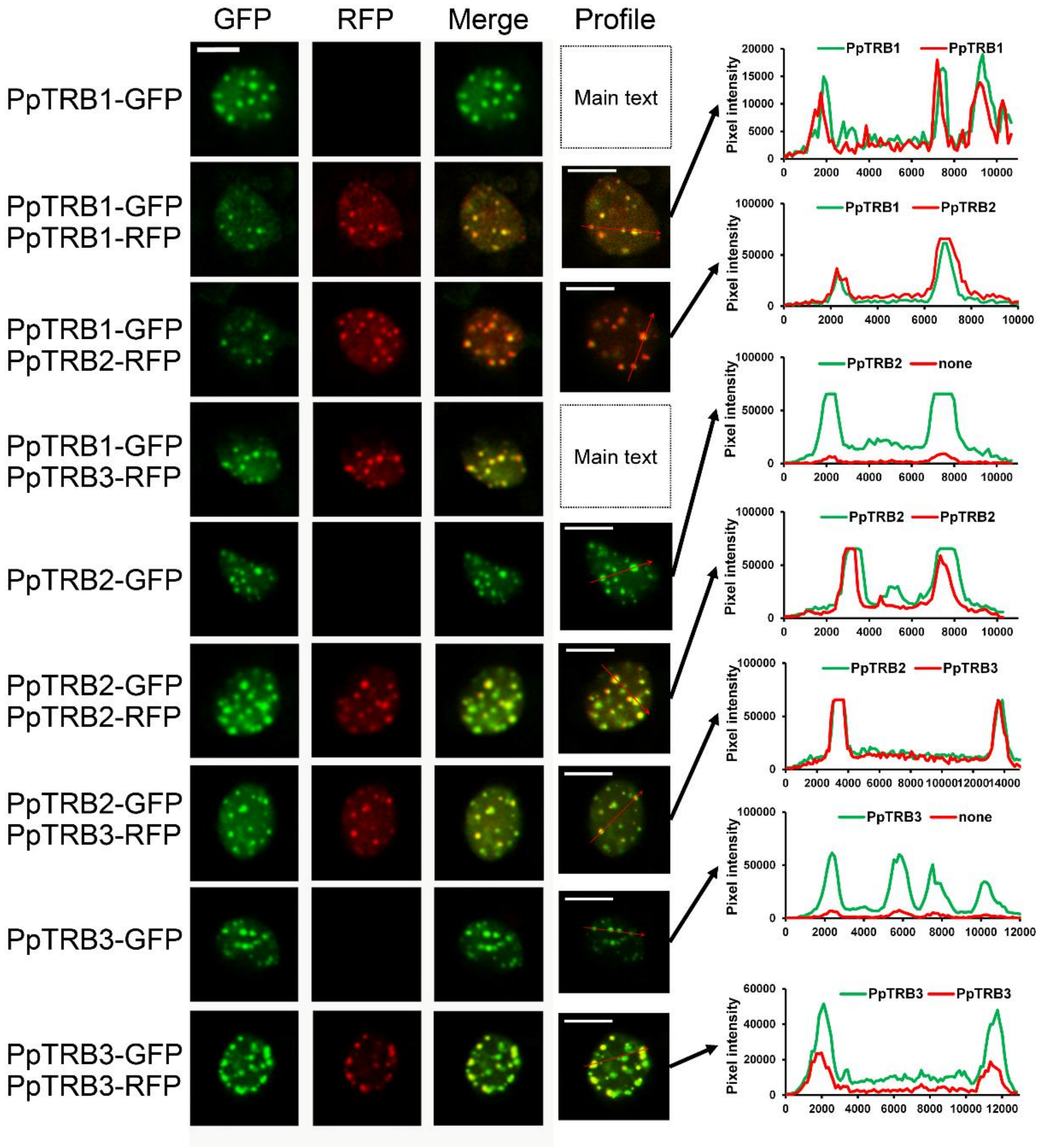


**Supplementary Figure 11. PpTRBs nuclear speckles locate at same spots in nucleoplasm.** Representative confocal microscopy images show the subnuclear localization of *PpTRB* proteins fused to GFP or RFP. Merged images reveal co-localization of *PpTRBs* at discrete nuclear foci. Scale bars = 5 µm. Intensity profiles corresponding to regions of interest (indicated by red arrows) confirm overlapping fluorescence signals, indicating co-localization of *PpTRBs* in nuclear speckles.

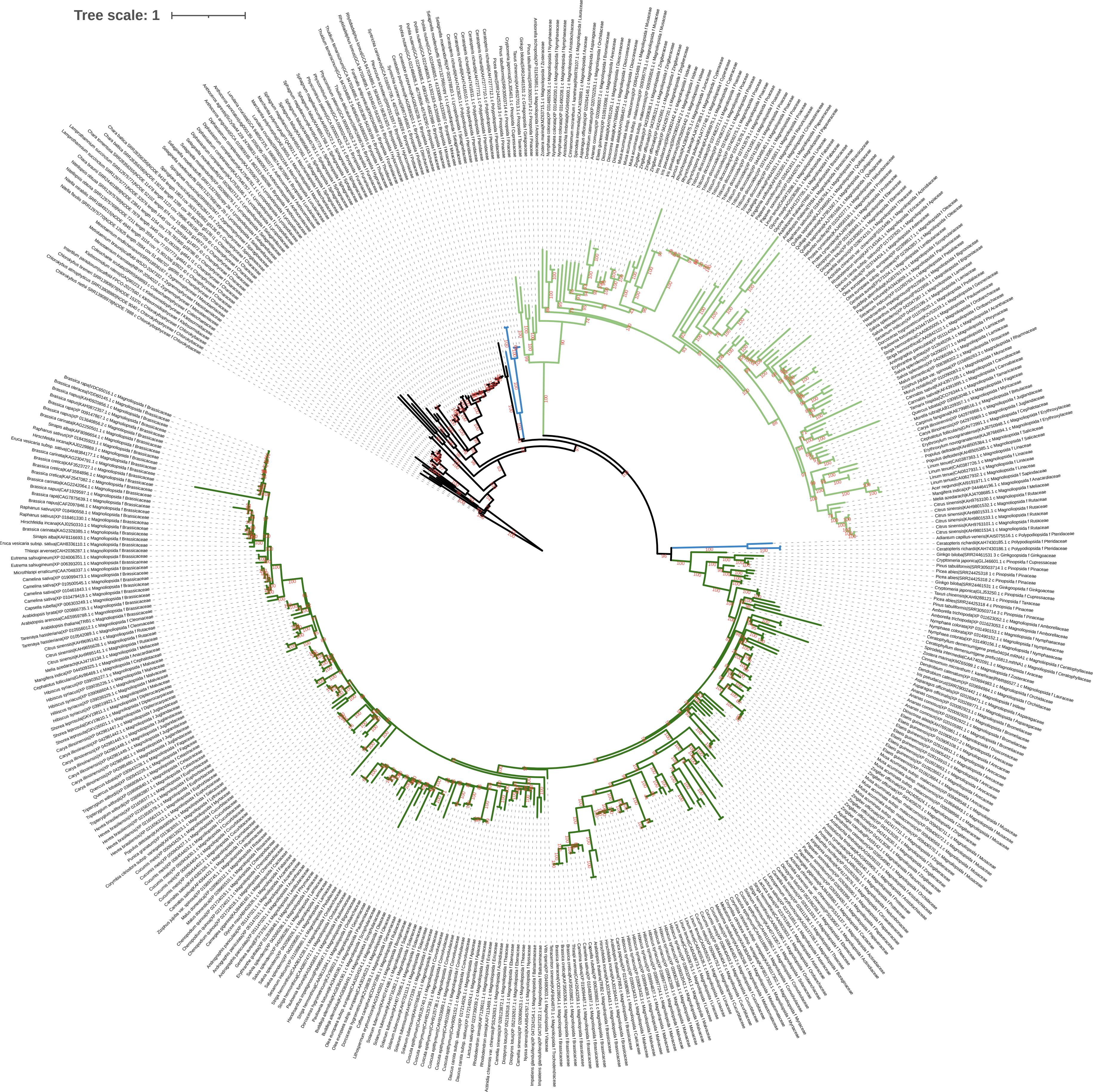
Tree scale: 1

**Supplementary Figure 12. Phylogenetic analysis of TRB proteins across streptophyte taxa.** Maximum likelihood (ML) phylogenetic trees were constructed to examine the evolutionary relationships of TRB proteins from various species, with accession numbers indicated for each sequence. Branch support was evaluated using ultrafast bootstrap approximation with 10,000 replicates; bootstrap values are shown below the branches. The trees were visualized using iTOL v6. Within Spermatophyta, TRB proteins segregate into two distinct clades, represented by light and dark green branches, highlighting their divergence in seed plants.
